# Molecular chaperone ability to inhibit amyloid-derived neurotoxicity, but not amorphous protein aggregation, depends on a conserved pH-sensitive Asp residue

**DOI:** 10.1101/2021.12.01.470723

**Authors:** Gefei Chen, Yuniesky Andrade-Talavera, Xueying Zhong, Sameer Hassan, Henrik Biverstal, Helen Poska, Axel Abelein, Axel Leppert, Nina Kronqvist, Anna Rising, Hans Hebert, Philip J.B. Koeck, André Fisahn, Jan Johansson

**Affiliations:** Department of Biosciences and Nutrition, Karolinska Institutet, 141 52 Huddinge, Sweden; Department of Neurobiology, Care Sciences and Society, Center for Alzheimer Research, Division of Neurogeriatrics, Neuronal Oscillations Laboratory, Karolinska Institutet, 171 77 Stockholm, Sweden; School of Engineering Sciences in Chemistry, Biotechnology and Health, Department of Biomedical Engineering and Health Systems, KTH Royal Institute of Technology, 141 52 Huddinge, Sweden; Department of Physical Organic Chemistry, Latvian Institute of Organic Synthesis, Riga LV-1006, Latvia; School of Natural Sciences and Health, Tallinn University, Tallinn, Estonia

**Author notes:** These authors contributed equally (Yuniesky Andrade-Talavera, Xueying Zhong, Sameer Hassan).

## Abstract

Proteins can self-assemble into amyloid fibrils or amorphous aggregates and thereby cause disease. Molecular chaperones can prevent both these types of protein aggregation, but the respective mechanisms are not fully understood. The BRICHOS domain constitutes a disease-associated small heat shock protein-like chaperone family, with activities against both amyloid toxicity and amorphous protein aggregation. Here, we show that the activity of two BRICHOS domain families against Alzheimer’s disease associated amyloid-β neurotoxicity to mouse hippocampi *in vitro* depends on a conserved aspartate residue, while the ability to suppress amorphous protein aggregation is unchanged by Asp to Asn mutations. The conserved Asp in its ionized state promotes structural flexibility of the BRICHOS domain and has a p*K*a value between pH 6.0–7.0, suggesting that chaperone effects against amyloid toxicity can be affected by physiological pH variations. Finally, the Asp is evolutionarily highly conserved in >3000 analysed BRICHOS domains but is replaced by Asn in some BRICHOS families and animal species, indicating independent evolution of molecular chaperone activities against amyloid fibril formation and non-fibrillar amorphous protein aggregation.

## Introduction

Proteins and peptides can self-assemble into highly ordered fibrillar structures as well as into smaller oligomers with less well-defined structures but apparently stronger ability to cause toxicity to cells. These phenomena are linked to so-called amyloid diseases, which encompass about forty severe human diseases including interstitial lung disease (ILD) and Alzheimer’s disease (AD)(1, 2). A multitude of proteins have been shown to form amyloid fibrils *in vitro* and generate cytotoxic species, yet they are not found in association with disease, which suggest that efficient chaperoning and/or other defence systems are in place(3, 4). Molecular chaperones prevent protein aggregation and cytotoxicity(5), and chaperonopathy might occur during *e.g.* ageing and by genetic mutations, leading to protein misfolding disorders(6–8). However, no clear correlation between the clinical phenotype and the severity of anti-aggregation defects of disease-causing molecular chaperone mutations has been shown for *e.g.,* DNAJB6b and HSPB1(9, 10).

Many exceptionally amyloidogenic polypeptides are expressed as proproteins which are subjected to proteolytic cleavage to release the amyloidogenic fragments(11–13). Some of these proproteins contain the small heat shock protein (sHSP)-like domain BRICHOS, initially found in integral membrane protein 2B (also called **Bri**2), **Cho**ndromodulin-1 and prosurfactant protein C (pro**S**P-C)(14, 15). The BRICHOS domain is supposed to promote the correct folding and prevent amyloid formation of the amyloid-prone region of its proprotein during biosynthesis(16–18), and is released by proteolysis(19–21). Mutations in the BRICHOS domain or in the proproteins are associated with different protein misfolding and amyloid diseases(16, 22, 23), but the underlying pathogenic mechanisms are largely unknown. The BRICHOS domain has been shown to have chaperone activities also in *trans* against amyloid fibril formation and toxicity of several peptides associated with human diseases(21, 24, 25) and has emerged as a model compound in studies of amyloid fibril formation(26–29). AD relevant amyloid-β with 42 residues (Aβ42) forms amyloid fibrils via several microscopic nucleation steps (*i.e.*, primary nucleation, elongation, and secondary nucleation), whereby Aβ42 oligomers are generated, which are supposed to be the main toxic species that drive AD development(27). The recombinant human (rh) BRICHOS domain from proSP-C is an efficient inhibitor of amyloid toxicity of Aβ42 *in vitro* in mouse hippocampal slice preparations, reducing the generation of toxic Aβ42 oligomers, but it is not very competent to reduce the overall amyloid fibril formation rate(24, 27). *In vivo* proSP-C BRICHOS has shown positive effects on locomotion and longevity when co-expressed with Aβ42 in a *Drosophila* fruit fly model(30, 31). The rh Bri2 BRICHOS, on the other hand, is efficient in inhibiting both Aβ42 amyloid fibril formation rate and toxicity by assembling into differently sized species, of which monomers potently prevent neuronal network toxicity of Aβ42, dimers strongly suppress Aβ fibril formation and large oligomers inhibit non-fibrillar protein aggregation(24, 30, 32, 33). Interestingly, for the BRICHOS domains studied so far, the ability to suppress amorphous protein aggregation, the canonical molecular chaperone function, is not coupled to the ability to prevent amyloid fibril formation or toxicity. ProSP-C BRICHOS essentially lacks activity against amorphous protein aggregation, while Bri2 and Bri3 BRICHOS inhibit both fibrillar and non-fibrillar aggregation(32, 34). We could recently explain the basis for this discrepancy by showing that the canonical chaperone activity to suppress amorphous protein aggregation is determined by specific loop segments in BRICHOS, while the activity against amyloid formation is independent of these segments (Chen et al, submitted for publication).

Recently, a Bri2 BRICHOS mutant (R221E), designed to stabilize the monomeric conformation, showed great potential in the prevention and treatment of AD by alleviating Aβ42 neurotoxicity rather than its overall fibrillization rate(33). A corresponding mutation in proSP-C BRICHOS (T187R) likewise generates monomers that bind to the smallest emerging Aβ42 oligomers(28). Surprisingly, T187N mutation of human proSP-C BRICHOS leads to ILD with amyloid deposits, while the T187R mutant is more efficient *in vitro* than the wildtype against amyloid fibril formation(28, 35). Thus, understanding the molecular mechanisms that regulate BRICHOS chaperone activities are of interest from a basic science point of view and for the development of treatments against amyloid diseases. In the BRICHOS superfamily, the secondary structure elements are highly conserved but the amino acid sequence conservation is low, and only one aspartic acid residue (Asp, D) and two cysteine residues that form a disulphide bridge were previously found to be strictly conserved, from analyses of a rather small number of sequences(14, 33). The physiological function of this Asp is unknown, yet two mutations of D105 in human proSP-C BRICHOS are linked to ILD(16, 20, 36). Herein, we analysed all BRICHOS proproteins deposited in sequence databases and studied experimentally how BRICHOS capacities against amyloid formation and toxicity, as well as non-fibrillar protein aggregation is affected by the conserved Asp. We found that the evolutionally conserved Asp is replaced by Asn in some BRICHOS families, is protonated at physiological pH and is essential for preventing amyloid toxicity but not amorphous aggregation.

## Results

### Evolutionary analyses of BRICHOS domains

As substantially more BRICHOS containing proteins have been described from genome sequencing since the initial analyses were performed, we undertook detailed phylogenetic and sequence analyses of all so far reported BRICHOS containing proteins. Amino acid sequences of 3 355 BRICHOS precursors were extracted from the SMART database, covering a broad distribution of 537 different metazoan species from worms to humans, grouped into eleven phyla (Fig. 1a **and** b). After sorting out repetitive and incomplete sequences, 2 019 BRICHOS sequences from 1 968 precursors were retained for further analysis. These BRICHOS domains were separated into thirteen groups, which include eleven previously known families (integral membrane protein 2A (ITM2A), ITM2B (also called Bri2), ITM2C (also called Bri3), group I, gastrokine-1 (GKN1), GKN2, GKN3, tenomodulin (TNMD), chondromodulin (CNMD, previously called LECTI), proSP-C, BRICHOS containing domain 5 (BRICD5) and two novel families (group II and antimicrobial peptide (AMP)) (Fig. 1c **centre and inner ring**, supplementary Fig. 1). The BRICHOS domains in the AMP group were recently found in marine animals to help antimicrobial peptides to fold correctly(37–39), while the BRICHOS domains in group II have unknown functions.

**Fig. 1.**
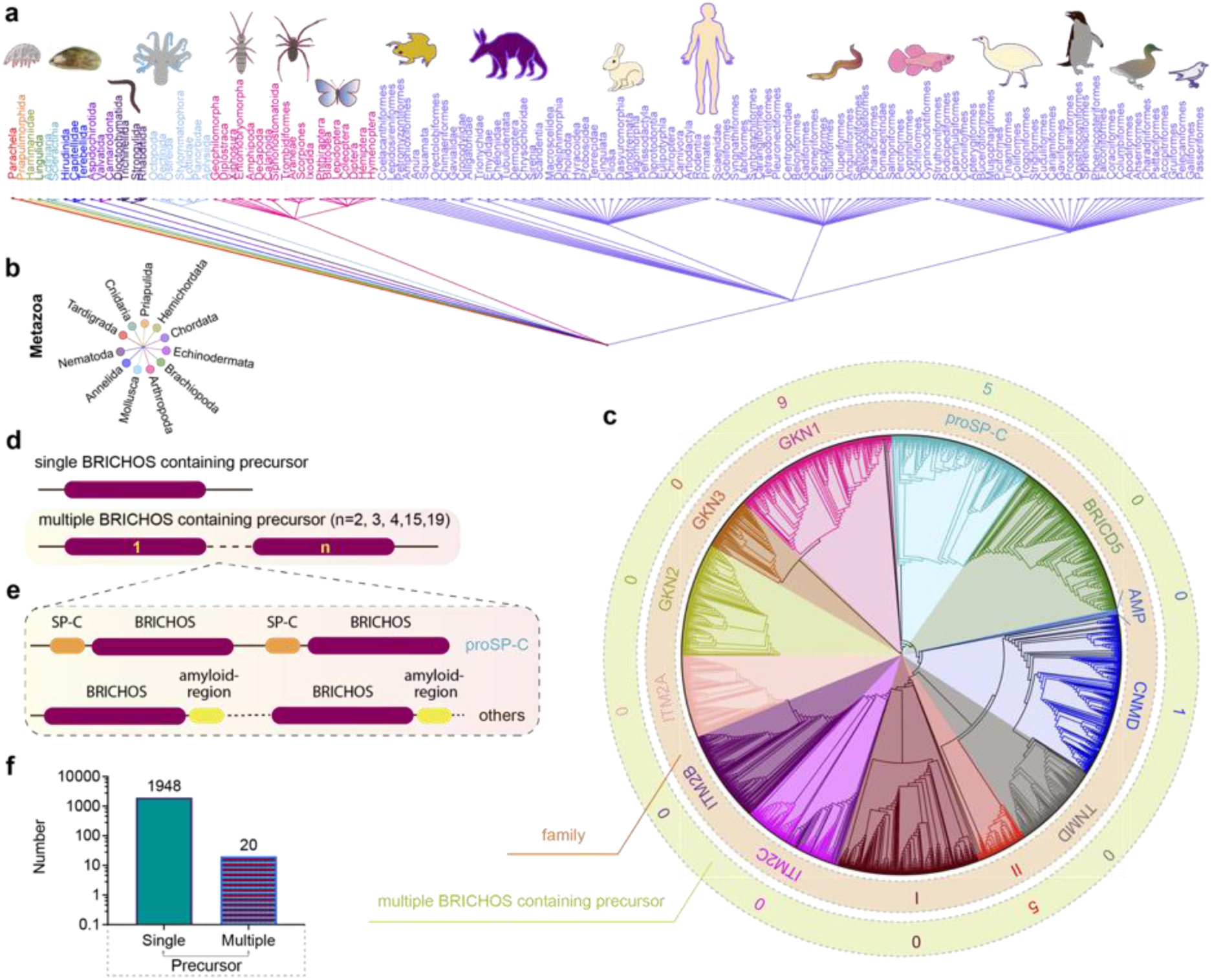
Evolutionary analyses of the BRICHOS domain. (**a**) The common taxonomy tree for all species containing BRICHOS domain precursors. The 3 355 BRICHOS precursors are distributed in a broad range of species, including worms, insects to human, and which belong to eleven metazoan phyla (**b**). (**c**) The selected 2 019 BRICHOS domains are grouped into thirteen families (inner ring), with bootstraps shown in Supplementary Figure 1. The outer ring shows the number of cases of occurrence of multiple BRICHOS domains in the respective families. (**d**) Architecture of proproteins containing single and up to nineteen multiple BRICHOS domains. The detailed architectures for each case are shown in Supplementary Figure 2. (**e**) the multiple BRICHOS domains showed there are a few BRICHOS containing proproteins that contain multiple BRICHOS domains along with representatives of their corresponding amyloid-prone regions. The true domain size is not proportional to the schematic bar. (**f**) The number of proproteins that contain single or multiple BRICHOS domains. 1 948 out of 1 968 proproteins contain a single BRICHOS domain while the others contain multiple BRICHOS domains.

Interestingly, analyses of the 1 968 precursors of the 2 019 BRICHOS domains showed that there are a few BRICHOS containing proproteins that contain multiple BRICHOS domains along with their corresponding amyloid-prone regions (Fig. 1d **and** e, Supplementary Fig. 2). Multiple BRICHOS domains were found in GKN1, proSP-C, CNMD and group II families (Fig. 1c **outer ring**). In total 99% (1948 out of 1968) of BRICHOS precursors contain a single BRICHOS domain, while 1% possess multiple BRICHOS domains (up to nineteen BRICHOS domains were found in one precursor) (Fig. 1f, Supplementary Fig. 2). The multiple BRICHOS domains within one precursor showed pairwise identities from 48% to 97% **(Supplementary Table 1**), indicating that they arose by duplications. Alignment of all compiled BRICHOS amino acid sequences showed that the Asp residue, which previously was found to be strictly conserved, did not show 100% conservation in neither the single nor the multiple BRICHOS domains (Fig. 2a **and** b, Supplementary Fig. 3 **and** 4). In the single BRICHOS domains, 97% (1853 out of 1908) contain the conserved Asp, while 2.7% have Asn, and the remaining 0.3% are distributed between Glu (E) and Ser (S) (Fig. 2c). Within the multiple BRICHOS domains, the percentage having Asp was markedly decreased to 31%, while instead 68% contain Asn, and one example with Glu was found (Fig. 2d). In the cases of multiple BRICHOS containing proproteins, 85% have either Asp or Asn, and 15% utilize both Asp and Asn (or Glu) (Fig. 2e, Supplementary Fig. 2). Among the BRICHOS domains derived from single BRICHOS precursors, the ones having Asn were found in GKN1, GKN3, ITM2A, and proSP-C BRICHOS families (Fig. 2a **and** f), and were mainly found in birds (Aves, 81%) with only 19% from Mammalia (Fig. 2f). An interesting observation is that bird surfactant lacks SP-C(40), so the BRICHOS containing proSP-C from birds likely have evolved other functions than to generate SP-C. Surprisingly, all the non-Asp BRICHOS domains from the multiple BRICHOS precursors were found in the GKN1 family and are all from birds (Fig. 2g).

**Fig. 2.**
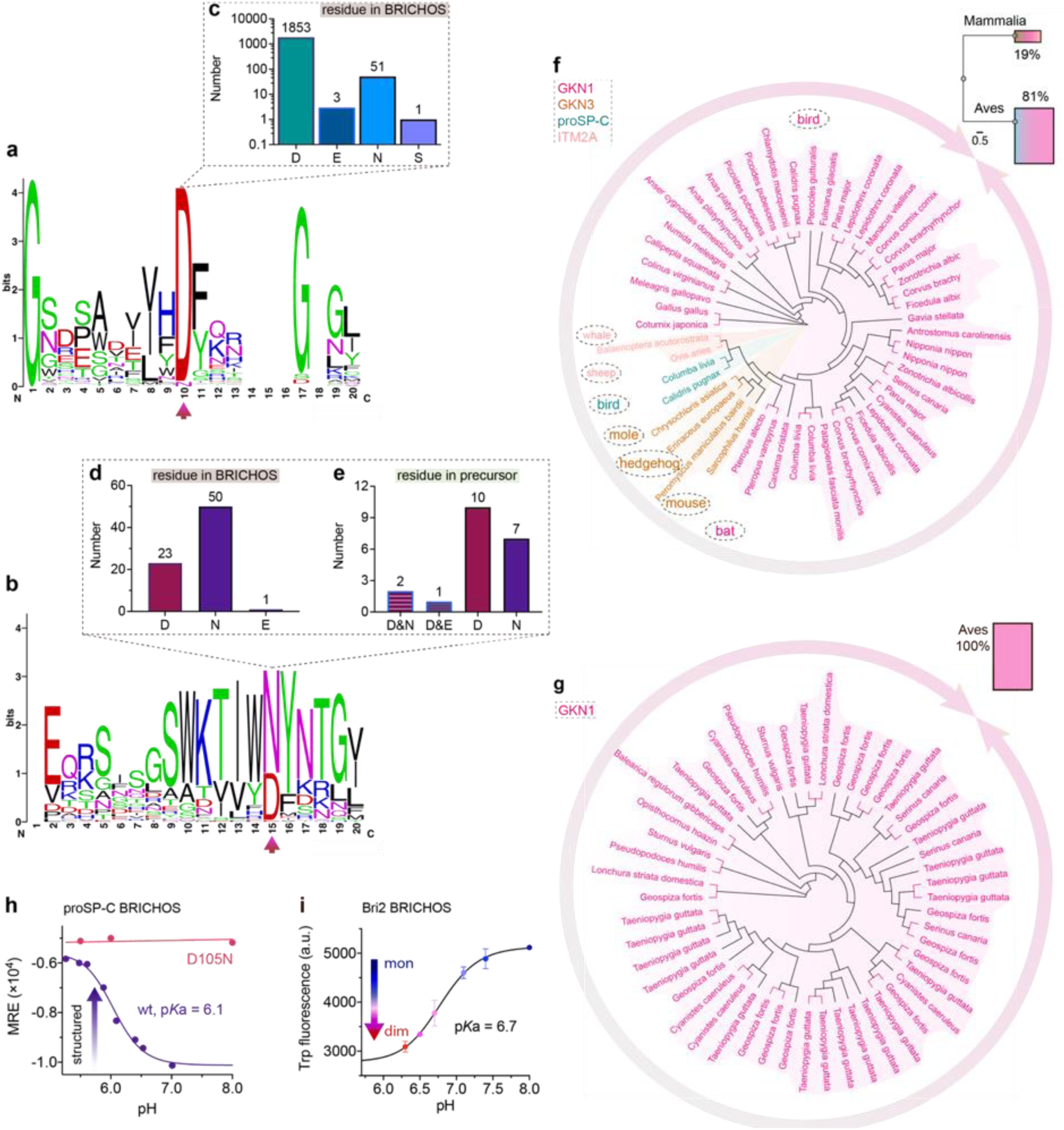
BRICHOS domain sequence analysis and species distribution. (**a**) WebLogo representation of amino acid sequence alignments of the 1908 single BRICHOS domains, surrounding the conserved Asp (red arrow). The height of the amino acid stack represents the sequence conservation, while the height of symbols within each stack indicates the relative frequency of each residue and the total height of the letters is given in bits. (**b**) WebLogo representation as in panel (**a**) of sequence alignments of the 74 multiple BRICHOS domains. (**c**) The residue distribution at the position of the conserved Asp in single BRICHOS domains. (**d**) The residue distribution at the position of the conserved Asp in multiple BRICHOS domains. (**e**) The consistency of Asp or Asn, or the combination of Asp and Asn (or Glu) in individual multiple BRICHOS containing precursor. (**f**) Family and species distributions of cases that contain Asn in single BRICHOS domains. (**g**) Family and species distributions of cases that contain Asn in multiple BRICHOS domains. (**h**) pH-dependent structural changes of rh proSP-C BRICHOS variants monitored by CD, the details are shown in Supplementary Figure 5b **and** c. (**i**) pH-dependent transition of rh Bri2 BRICHOS monomers to dimers measured by Trp fluorescence, the details are shown in Supplementary Figure 9b **and** c.

### Ionization and protonation of the Aspartate within a physiological pH range

The results from the evolutionary studies prompted us to mutate the conserved Asp to Asn in human BRICHOS domains from proSP-C (Asp105) and Bri2 (Asp148), which are the most studied BRICHOS domains.

Size exclusion chromatography (SEC) showed that both wildtype and D105N rh proSP-C BRICHOS adopt trimeric conformation, but the mutant formed more dimers and less monomers than the wildtype protein (Supplementary Fig. 5a). Circular dichroism (CD) spectroscopy showed rh proSP-C BRICHOS D105N is overall more structured than the wildtype at neutral pH as judged from a shift from the minimum around 202 nm to around 207 nm, while at acidic pH the CD spectra are virtually identical (Supplementary Fig. 5b **and** c). To see whether wildtype rh proSP-C BRICHOS titrates with an elevated p*K*a value (compared to p*K*a of about 4.5 for free Asp in solution), we followed the CD signal at 204 nm of rh proSP-C BRICHOS between pH 7.0 and 5.0, which gave an apparent p*K*a value of the transition of about 6.1 (Fig. 2h, Supplementary Fig. 5b). Notably, the CD spectra of rh proSP-C BRICHOS D105N did not shift substantially between pH 5.5 and 8.0 (Fig. 2h, Supplementary Fig. 5c), suggesting that protonation of D105 is a main determinant of the observed conformational shift, and it may be the sole titrating residue in this pH range. To confirm the pH-dependent conformational changes, we ran NMR spectroscopy on wildtype rh proSP-C BRICHOS labelled with ^15^N, ^13^C and ^2^H at different pHs. The comparison of the ^15^N-HSQC spectra at pH 5.5 and 7.2 showed clear chemical shift changes, supporting that rh proSP-C BRICHOS undergoes structural changes in this pH range (Supplementary Fig. 5d).

Rh wildtype NT*-Bri2 BRICHOS(33) and rh NT*-Bri2 BRICHOS D148N showed similar elution profiles on SEC (Supplementary Fig. 5e). After removal of the NT* solubility tag, rh Bri2 BRICHOS D148N formed similar disulphide-dependent assembly states as the wildtype protein (Supplementary Fig. 5f–h). However, the rh Bri2 BRICHOS D148N dimers are exclusively disulphide-linked (Supplementary Fig. 5h, **Figure 5-figure supplement 5-source data**), while the wildtype dimers are a mixture of disulphide-linked and non-covalent species(32). Moreover, rh Bri2 BRICHOS D148N generated from the rh NT*-Bri2 BRICHOS D148N monomer fraction did not elute on SEC as a monomer but as a dimer, while both the corresponding oligomers and dimers shared similar elution profiles as the corresponding wildtype states (Supplementary Fig. 5f **and** g). This indicates that the rh Bri2 BRICHOS D148N monomers are not stable but form non-covalent dimers. Rh Bri2 BRICHOS D148N oligomers and dimers shared identical CD spectra as the corresponding wildtype species, whereas the initially monomeric rh Bri2 BRICHOS D148N showed a somewhat different CD spectrum compared to the wildtype monomers (Supplementary Fig. 5i–l), but which can be superimposed on the wildtype dimer spectrum (Supplementary Fig. 5l). This suggests the rh Bri2 BRICHOS D148N monomer mainly assembles into a dimer at the concentration used for CD spectroscopy. As analysed by SEC, at one µmol L^-1^, rh wildtype Bri2 BRICHOS monomer migrated solely as a monomer, and with increasing concentration, dimers were progressively formed (Supplementary Fig. 6a, Supplementary Fig. 7). However, an equilibrium between dimers and monomers were seen even at 200 µmol L^-1^ (Supplementary Fig. 6a, Supplementary Fig. 7e). In contrast, already at low concentrations (0.5 µmol L^-1^) rh Bri2 BRICHOS D148N showed an equilibrium between dimers and monomers, approximately at a ratio of 1:1 (Supplementary Fig. 6b, Supplementary Fig. 8a). With progressively increased concentrations rh Bri2 BRICHOS D148N monomers transformed to dimers and at 100 µmol L^-1^ all proteins migrated as dimers (Supplementary Fig. 6b, Supplementary Fig. 8f). To get further details on how rh Bri2 BRICHOS undergoes monomer-dimer transition, we analysed SEC profiles at different pHs. At pH 7.0 to 8.0 both rh Bri2 BRICHOS D148N and wildtype monomers showed similar elution volumes (Supplementary Fig. 6c **and** d). However, at pH 6.0 rh wildtype Bri2 BRICHOS elutes as a dimer, whereas the D148N mutant did not show any significant difference compared to the elution volume at pH 7.0 and 8.0 (Supplementary Fig. 6c **and** d).

This suggests that at pH between 7.0 and 6.0 Asp148 can be protonated with subsequent shift of the dimer-monomer equilibrium towards dimers. To corroborate the supposition that Asp148 titrates between pH 6.0 and 7.0 we turned to a rh Bri2 BRICHOS variant with Thr206 replaced by Trp (T206W) that displayed an identical oligomerization profile as the rh wildtype Bri2 BRICHOS (Supplementary Fig. 9a), but a different Trp fluorescence profile in the monomeric state compared to the dimeric state (Supplementary Fig. 9b). In agreement with SEC data, titration of the rh Bri2 BRICHOS Trp mutant showed a fluorescence evolution (monomer/dimer transition) with a p*K*a of 6.7 (Fig. 2i, Supplementary Fig. 9c).

These observations indicate that the D148N mutation in rh Bri2 BRICHOS promotes monomer to dimer conversion, and mimics pH-induced dimerization of the wildtype monomers with an apparent p*K*a of 6.7, but it does not significantly change the assembly of dimers into larger oligomers. Analogously, the mutation D105N in rh proSP-C BRICHOS results in a more compact conformation and possibly more complete assembly into trimers, and the wildtype rh proSP-C BRICHOS conformational shift by CD titrates with an apparent p*K*a of 6.1.

### The conserved Asp of BRICHOS is essential for the capacity to prevent Aβ42-induced neurotoxicity

To investigate the importance for the ability to alleviate amyloid-induced neurotoxicity, we tested the efficacies of Asp to Asn mutated vs. wildtype BRICHOS in preventing Aβ42-induced reduction of γ oscillations in mouse hippocampal slices. γ oscillations correlate with learning, memory, cognition and other higher processes in the brain(41, 42) and progressive cognitive decline observed in AD goes hand-in-hand with a progressive decrease of γ oscillations(43–45). The BRICHOS domains from human proSP-C and Bri2 can efficiently prevent Aβ42-induced decline in hippocampal γ oscillations(27, 30, 32, 33, 46). In addition, rh Bri2 BRICHOS rescues γ oscillations and neuronal network dynamics from Aβ42-induced impairment at hippocampal CA3 area *ex vivo*(47). We recorded γ oscillations in hippocampal slices from wildtype C57BL/6 mice preincubated for 15 min either with 50 nmol L^-1^ Aβ42 alone, or co-incubated with 100 nmol L^-1^ rh wildtype or D105N proSP-C BRICHOS (Fig. 3a). γ oscillations were elicited by application of 100 nmol L^-1^ kainic acid (KA) and allowed to stabilize for 30 min prior to any recordings. As previously observed(27, 30, 46), 100 nmol L^-1^ wildtype proSP-C BRICHOS prevented Aβ42-induced degradation of γ oscillations, which were remained at control levels (Fig. 3b **and** c, control: 1.7 ± 0.16 × 10^-8^ V^2^ Hz^-1^, n= 20; Aβ42: 0.34 ± 0.06 × 10^-8^ V^2^ Hz^-1^, n= 14, *p*< 0.0001 vs. control; + rh wildtype proSP-C BRICHOS: 1.41 ± 0.24 × 10^-8^ V^2^ Hz^-1^, n= 9) (Supplementary Table 2, **Figure 3-source data**). By contrast, rh proSP-C BRICHOS D105N showed a complete loss of the preventative efficacy at the same concentration (Fig. 3b **and** c, rh proSP-C BRICHOS D105N: 0.36 ± 0.09 × 10^-08^ V^2^ Hz^-1^, n= 9, *p*= 0.0202 vs. rh wildtype proSP-C BRICHOS, *p*= 0.0002 vs. control, *p*> 0.9999 vs. Aβ42) (**Supplementary Table 2**, **Figure 3-source data**). We have previously observed that the rh Bri2 BRICHOS monomer is most efficient at preventing Aβ42 induced degradation of γ oscillations(32, 33). We therefore tested whether the D148N mutation could affect the efficacy of rh Bri2 BRICHOS monomers. Rh Bri2 BRICHOS D148N monomers (50 nmol L^-1^) showed reduced tendency of potency to prevent Aβ42-induced neurotoxicity compared to wildtype monomers (50 nmol L^-1^), but did not completely lose preventative efficacy, as observed for rh proSP-C BRICHOS D105N (Fig. 3d **and** e, rh wildtype Bri2 BRICHOS monomers: 1.35 ± 0.29 × 10^-8^ V^2^ Hz^-1^, n= 8, *p*> 0.9999 vs. control; rh Bri2-BRICHOS D148N monomers: 0.7 ± 0.14 × 10^-8^ V^2^ Hz^-1^, n= 11, *p*= 0.3494 vs. rh wildtype Bri2 BRICHOS monomers, *p*= 0.003 vs. control, *p*= 0.6991 vs. Aβ42) (**Supplementary Table 2**, **Figure 3-source data**). These results suggest that the phylogenetically conserved Aspartate plays a crucial role in maintaining the anti-amyloid neurotoxicity capacity of BRICHOS.

**Fig. 3.**
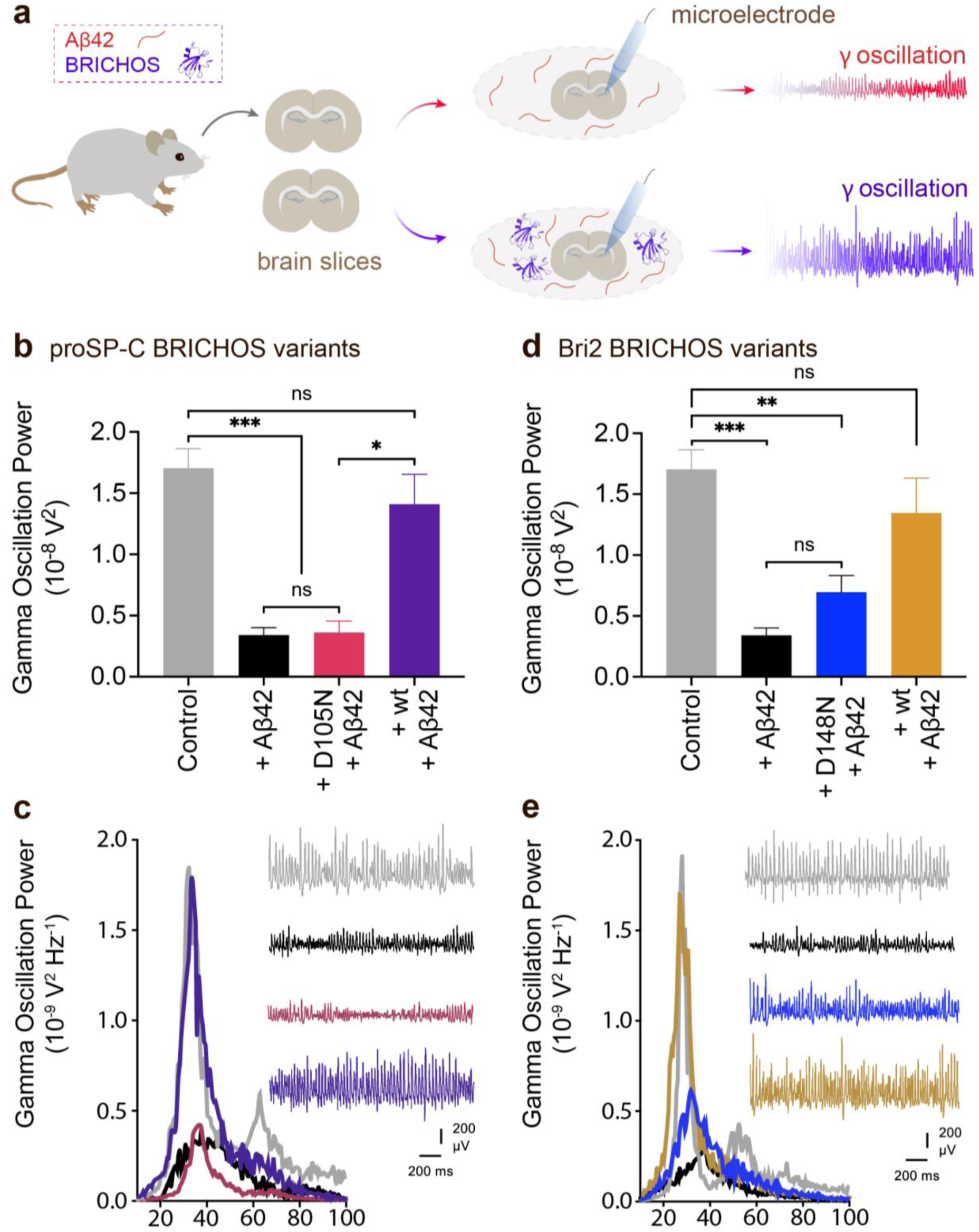
Asp to Asn mutation reduces rh proSP-C BRICHOS and rh Bri2 BRICHOS capacity against Aβ42 neurotoxicity. (**a**) Schematic diagram of electrophysiological recordings. The hippocampal slices from wildtype C57BL/6 mice were preincubated either with 50 nmol L^-1^ Aβ42 alone or co-incubated with 100 nmol L^-1^ rh BRICHOS, and γ oscillations were then recorded in the CA3 area. (**b**) Summary plot of normalized γ oscillation power under control conditions (gray, n=20), after 15 min incubation with 50 nmol L^-1^ Aβ42 (black, n=14), 50 nmol L^-1^ Aβ42 + 100 nmol L^-1^ D105N (red, n=9) or wildtype (purple, n=9) rh proSP-C BRICHOS. Example traces and example power spectra are shown with the same color coding in (**c**). (**d**) Summary plot of γ oscillation power under control conditions (gray, n=20), after 15 min incubation with 50 nmol L^-1^ Aβ42 (black, n=14), 50 nmol L^-1^ Aβ42 + 100 nmol L^-1^ D148N monomeric (blue, n=11) or wildtype monomeric (yellow, n=8) rh Bri2 BRICHOS. Example traces and example power spectra are shown with the same color coding in (**e**). The data are reported as means ± standard errors of the means. ns, no significant difference, \**p* < 0.05, \*\**p* < 0.01, \*\*\**p* < 0.001.

### The conserved Asp of BRICHOS is not important for canonical chaperone activity against amorphous protein aggregation or Bri2 BRICHOS oligomer structure

The rh wildtype Bri2 BRICHOS oligomers are efficient molecular chaperones against non-fibrillar protein aggregation, evaluated by thermo-induced citrate synthase aggregation as a model(32). This was found to be true also for the rh Bri2 BRICHOS D148N oligomers (Fig. 5a and b). Similarly, the D105N mutation did not show any effects on the capacity of rh proSP-C BRICHOS trimer against amorphous protein aggregation, *i.e.*, both the wildtype and D105N mutant are essentially inactive as regards canonical chaperone activity (Supplementary Fig. 10a). This shows a strong difference in the importance of the conserved Asp in BRICHOS for activity against amyloid toxicity compared to activity against amorphous protein aggregation.

Although the D148N mutation changed the balance between monomers and dimers of rh Bri2 BRICHOS, the mutant still formed large oligomeric species with a similar secondary structure as wildtype oligomers (Supplementary Fig. 5e, g **and** i). We isolated rh Bri2 BRICHOS D148N oligomers by SEC, recorded transmission electron microscopy (TEM) micrographs and calculated 3D reconstructions (Fig. 4c **and** d, Supplementary Fig. 10b **and** c, **Figure 4-source data**). The micrographs and 2D class averages revealed mostly homogenous large assemblies with two-fold symmetry (Fig. 4c). As a result, an EM map was reconstructed with D2 symmetry from 10 223 manually extracted single particles (Fig. 4d, **Figure 4-source data**). The rh Bri2 BRICHOS D148N oligomer 3D model and the one from the wildtype oligomers(32) overlap and share overall similar shape and volume (Fig. 4e, **Figure 4-source data**), suggesting that the overall structure of the large Bri2 BRICHOS oligomers is not noticeably changed when Asp148 is mutated to Asn.

**Fig. 4.**
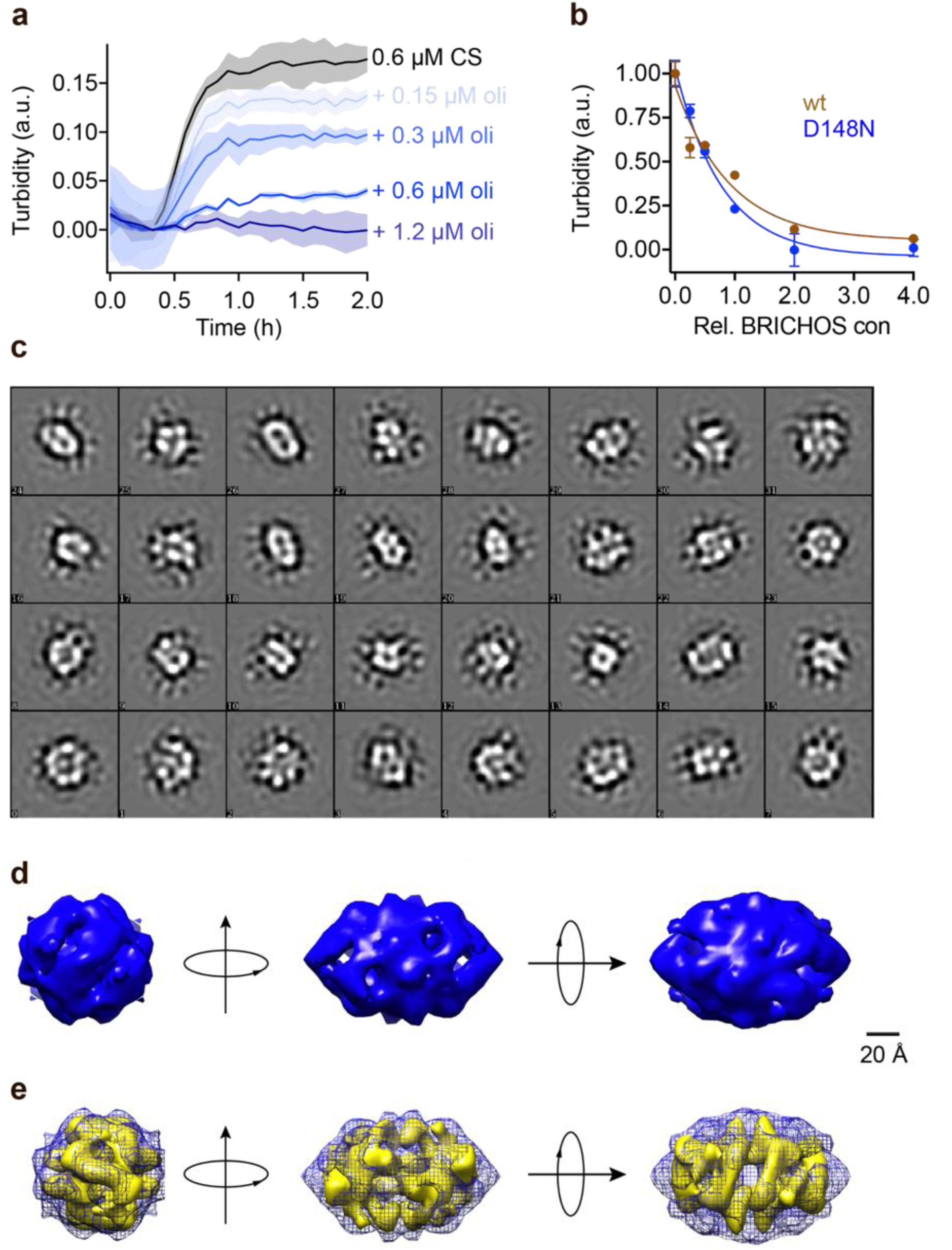
Rh Bri2 BRICHOS D148N oligomer EM map and comparison with the wildtype oligomer 3D EM density map. (**a**) Kinetics of aggregation of 600 nmol L^-1^ citrate synthase (CS) at 45 °C alone (black), in the presence of 0.15, 0.3, 0.6 and 1.2 µmol L^-1^ rh Bri2 BRICHOS D148N oligomer. The different concentrations of the oligomers are shown with a blue gradient, which are labelled on the right of the traces. (**b**) Effects of rh Bri2 BRICHOS D148N and wildtype oligomers (data from ref ^(32)^) on CS aggregation at different molar ratios (referred to monomeric subunits) of BRICHOS:CS. The data are presented as means ± standard deviations. (**c**) 2D classes of rh Bri2 BRICHOS D148N oligomers. Most class averages show an approximate 2-fold symmetry. (**d**) The three viewing directions are along the three different 2-fold axes (upper panel). The map of rh Bri2 BRICHOS D148N oligomer was based on 10 223 particles manually extracted from images recorded on a CCD detector with ×85 200 magnification and the voxel size of the map is 2.464 Å. The lower panel shows the 3D density map of rh Bri2 BRICHOS D148N oligomer with D2 symmetry (blue) and the volume fit in rh wildtype Bri2 BRICHOS oligomer (yellow, EMDB: 3918).

### Molecular mechanisms underlying the importance of the conserved Asp of BRICHOS for ability to suppress Aβ42 neurotoxicity

To further explore whether the diminished BRICHOS capacity against amyloid neurotoxicity caused by Asp to Asn mutation correlates with the activity of suppressing macroscopic amyloid fibril formation, we used thioflavin T (ThT)(48) fluorescence to monitor the kinetics of Aβ42 fibril formation in the absence and presence of different concentrations of rh Bri2 or proSP-C BRICHOS (Fig. 5). Both the rh wildtype proSP-C BRICHOS and the D105N mutant showed dose-dependent progressive reduction of Aβ42 fibril formation at substoichiometric concentrations, and the aggregation kinetics follows a typical sigmoidal behavior (Fig. 5a **and** d–f, Supplementary Fig. 11a-c). The Aβ42 fibrillization half time, *τ*_1/2_, increases with increasing rh proSP-C BRICHOS concentration (Fig. 5b), while the maximum rate of Aβ42 aggregation, *r_max_*, shows a mono-exponential decline (Fig. 5c). Interestingly, rh proSP-C BRICHOS D105N showed remarkably improved inhibition on Aβ42 fibril formation compared to the wildtype, manifested with both *τ*_1/2_ and *r_max_* (Fig. 5a–c), which is opposite compared to the effects on the capacity to prevent Aβ42-induced neurotoxicity (Fig. 3). On the other hand, similarly, the rh wildtype Bri2 BRICHOS monomers showed similar effects on *τ*_1/2_ and *r_max_* of Aβ42 fibrillization as reported (Fig. 5g–i)(32). The rh Bri2 BRICHOS D148N oligomers, dimers and monomers also presented dose-dependent inhibition on Aβ42 fibril formation with typical sigmoidal behavior (Fig. 5g–i, Supplementary Fig. 12). Along the same lines, compared to wildtype monomers, the D148N monomers were significantly more efficient in inhibiting Aβ42 fibril formation (Fig. 5g–i), suggesting the Asp to Asn mutation pronouncedly enhances BRICHOS activity against overall fibril formation. The D148N monomers and dimers showed very similar inhibitory effects on Aβ42 fibrillization (Supplementary Fig. 12a **and** b), which likely can be explained by that D148N monomers and dimers are in equilibrium (Supplementary Fig. 6b **and** 8).

**Fig. 5.**
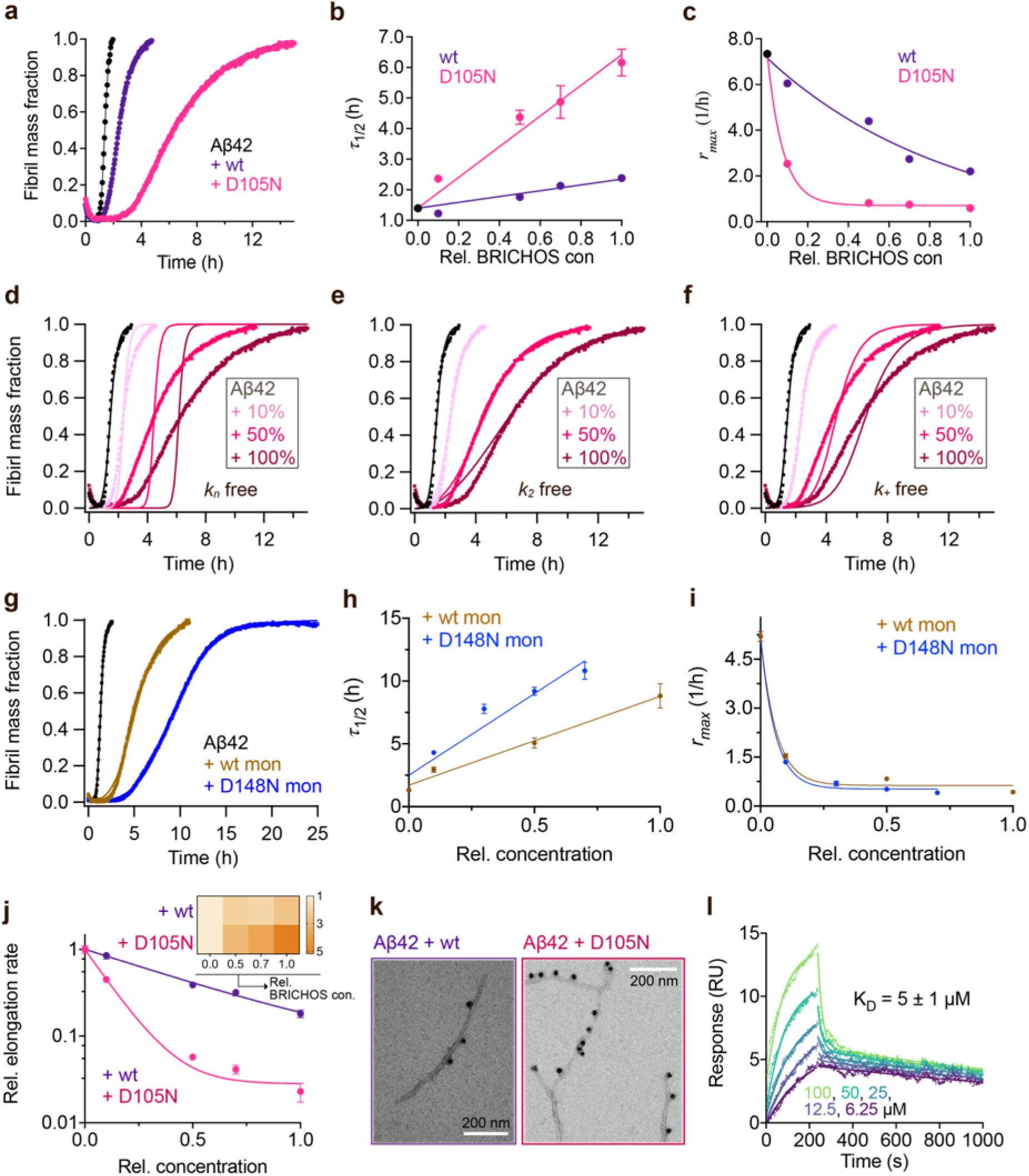
Asp to Asn mutation changes rh proSP-C BRICHOS activity against Aβ42 fibril formation. (**a**) Activity comparison of 100% rh wildtype proSP-C BRICHOS (purple) and rh proSP-C BRICHOS D105N (red) against 3 µmol L^-1^ Aβ42 (black). Individual fits with combined rate constants 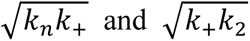 as free fitting parameters of normalized and averaged aggregation traces (dots) are shown as solid lines. Values for *τ_1/2_* (**b**) and *r_max_* (**c**) extracted from sigmoidal fitting of Aβ42 aggregation traces in the presence of different concentrations of rh wildtype proSP-C BRICHOS (purple) or rh proSP-C BRICHOS D105N (red) as shown in (**d-f**) and (Supplementary Figure 10a–c). (**d–e**) Aggregation kinetics of 3 µmol L^-1^ Aβ42 in the presence of rh proSP-C BRICHOS D105N at concentrations: 0 (black), 10 (light red), 50 (red), or 100% (dark red) molar percentage referred to monomeric subunits relative to Aβ42 monomer. The global fits (solid lines) of the aggregation traces (crosses) were constrained such that only one single rate constant is the free fitting parameter, indicated in each panel. χ^2^ values describing the quality of the fits: 18 for *k_n_* free, 0.8 for *k*_2_ free and 3.2 for *k*_+_ free. (**g**) Comparison of 50% rh wildtype Bri2 BRICHOS (yellow) and rh Bri2 BRICHOS D148N (blue) activities against 3 µmol L^-1^ Aβ42 (black). The solid lines are from individual fits with combined rate constants 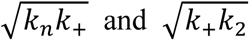 as free fitting parameters of normalized and averaged aggregation traces (dots). Values for *τ_1/2_* (**h**) and *r_max_* (**i**) extracted from the sigmoidal fitting of Aβ42 aggregation traces in the presence of different concentrations of rh wildtype Bri2 BRICHOS and the D148N mutant as shown in (Supplementary Figure 10c–e). (**j**) Elongation rates (*k_+_*) determined from highly seeded aggregation kinetics in Supplementary Figure 10d **and** e. The inset shows the amount of toxic Aβ42 oligomers calculated with the relative elongation rates (*k_+_*) and secondary nucleation rates (*k*_2_) for either rh wildtype proSP-C BRICHOS or the D105N mutant. (**k**) Immuno-EM of BRICHOS bound to Aβ42 fibrils. Aβ42 was incubated with and without 100% molar ratio of rh wildtype proSP-C BRICHOS and the D105N mutant, respectively, overnight at 37°C. The samples were treated with a polyclonal antibody against human proSP-C and a gold-labelled secondary antibody, and characterized by TEM. The scale bars are 200 nm. (**l**) SPR sensorgrams of different concentrations (*i.e.,* 6.25, 12.5, 25, 50, and 100 µmol L^-1^) of rh proSP-C BRICHOS D105N interacting with immobilized Aβ42 monomers. The data were globally fitted for the association and disassociation phases, respectively, and the apparent *K*_D_ was calculated.

To find out the molecular mechanisms underlying how the Asp to Asn mutation decreases BRICHOS capacity against Aβ42 neurotoxicity but enhances its inhibitory activity against overall Aβ42 fibril formation, we investigated the effects of both rh wildtype BRICHOS and the corresponding Asp to Asn mutants on Aβ42 amyloid fibril formation and related microscopic events. Aβ42 fibrillization kinetics are described by a set of microscopic rate constants, *i.e.*, primary (*k_n_*) and surface-catalyzed secondary nucleation (*k*_2_) as well as elongation (*k_+_*)(49), and perturbations of the individual microscopic rates are most relevant since their relative contributions determine the generation of nucleation units, which might be linked to the neurotoxic Aβ42 oligomeric species(27, 50). To evaluate the effects on individual microscopic processes, we performed global fits of the kinetics data sets at constant Aβ42 and different BRICHOS concentrations for both rh wildtype proSP-C and the D105N mutant, where the fits were constrained such that only one single rate constant, *i.e., k_n_*, *k*_2_ or *k_+_*, is the sole fitting parameter (Fig. 5d-f, Supplementary Fig. 11a–c). As previously described(27), the rh wildtype proSP-C BRICHOS mainly interfered with the secondary nucleation, indicated by the perfect fits when *k*_2_ was the sole global fitting rate constant (Supplementary Fig. 11a–c). Also, the secondary nucleation rate *k*_2_ as the sole fitting parameter gave the best fits for the Aβ42 fibrillization kinetics with rh proSP-C BRICHOS D105N (Fig. 5d-e), but with worse quality compared to the wildtype, especially for the start of the aggregation traces. This suggests that a complex microscopic mechanism is present, which might also include fibril-end elongation (*k_+_*) and/or primary nucleation (*k_n_*) in addition to secondary nucleation (*k*_2_). To further study whether the fibril-end elongation process or the primary nucleation are affected in the presence of rh proSP-C BRICHOS D105N and wildtype forms, we determined aggregation kinetics in the presence of a high initial fibril seed concentration(49) and surface plasmon resonance (SPR) analysis. With seeding, the fibrillization traces typically follow a concave aggregation behavior (Supplementary Fig. 11d **and** e), where the initial slope is directly proportional to the elongation rate *k_+_*. These seeding experiments revealed that rh proSP-C BRICHOS D105N decreases the elongation in a dose-dependent manner, and already at low concentrations fibril-end elongation is noticeably retarded (Fig. 5j, Supplementary Fig. 11e). The rh wildtype proSP-C BRICHOS, in contrast, showed only slight effects on fibril-end elongation (Fig. 5j, Supplementary Fig. 11d), which is further supported by immuno-transmission electron microscopy (immuno-EM) images where both rh proSP-C BRICHOS D105N and the wildtype attach along the Aβ42 fibril surface, however, the fibril ends are apparently mainly blocked by the D105N mutant (Fig. 5k). Additionally, SPR analyses of Aβ42 monomers immobilized on a sensor chip indicated that rh wildtype proSP-C BRICHOS showed weak binding to the immobilized Aβ42 monomers in line with previous reports(27, 28). The D105N mutation significantly enhanced the BRICHOS-Aβ42 monomer interactions (Supplementary Fig. 11f) with an apparent affinity value *K*_D_ around 5 µmol L^-1^ (Fig. 5l) from global kinetics fits, and under steady-state conditions a *K*_D_ value of around 25 µmol L^-1^ was obtained (Supplementary Fig. 11g). Interfering with primary nucleation (*k_n_*) delays Aβ42 fibril formation without changing the total number of oligomers generated, suppressing the secondary nucleation (*k*_2_) efficiently prevents the generation of oligomers, while blocking elongation (*k*_+_) significantly increases the number of oligomers formed(27). Of relevance for the results on Aβ42 neurotoxicity of wildtype vs D105N proSP-C BRICHOS (Fig. 3), we found that in the presence of equimolar ratio of rh proSP-C BRICHOS D105N, the Aβ42 fibrillization reaction generated approximately 5.2 times more oligomers than from Aβ42 alone, by significantly suppressing the fibril end elongation process, while the wildtype only showed slight effects (Fig. 5j **insets**). These observations show that both rh proSP-C BRICHOS D105N and the wildtype protein reduce Aβ42 fibrillization by blocking the surface-catalyzed secondary nucleation, while fibril-end elongation and possibility primary nucleation are substantially affected only by the D105N mutant. These results offer a molecular explanation to the observed loss of inhibitory effects of the rh proSP-C BRICHOS mutant on neurotoxic Aβ42 oligomer generation.

To further study the mechanism underlying the reshaped interference of BRICHOS with Aβ42 monomer and fibrils, we used the non-polar fluorescent dye bis-ANS to probe the exposure of hydrophobic areas. Bis-ANS shows a blue shift of the emission maximum and increased emission intensity upon binding to exposed hydrophobic protein surfaces, which has been applied to rh wildtype Bri2 and proSP-C BRICHOS(32, 51, 52). When incubated with bis-ANS, rh wildtype proSP-C BRICHOS gave an increase of emission intensity compared to bis-ANS in buffer, and a blue shift of the emission maximum from about 520 nm to 495 nm (Supplementary Fig. 11h). Notably, the blue shift and the intensity increase for the D105N mutant at pH 8.0 are more pronounced compared to the wildtype protein (Supplementary Fig. 11h), showing that D105N results in more exposed hydrophobic areas, which may be important for Aβ42 fibril-end binding(32, 33) and monomer binding. Interestingly, when incubating bisANS with rh wildtype proSP-C BRICHOS at pH 6.0 an identical fluorescence spectrum as for the D105N mutant at pH 8.0 was observed (Supplementary Fig. 11h), indicating that the D105N mutation gives similar effects as protonating Asp105. For Bri2 BRICHOS, dimers were found previously to be most efficient in preventing Aβ42 overall fibril formation compared to the monomers and oligomers, while the monomers are most potent in preventing Aβ42 induced disruption of neuronal network activity(32, 33). The microscopic mechanisms of rh Bri2 BRICHOS D148N species were similar to the wildtype species, both the secondary nucleation and elongation of Aβ42 were affected (Supplementary Fig. 12c–k). However, the equilibrium between monomer and dimer was modified by the Asp to Asn mutant towards the dimer (Supplementary Fig. 6a **and** b, Supplementary Fig. 7 **and** 8), thus reducing the monomer that is predominantly active against Aβ42-induced neurotoxicity but increasing the dimer fraction, which is most active against Aβ42 fibril formation.

## Discussion

In this work, we show that the capacity of sHSP-like chaperone domain BRICHOS to inhibit amyloid-associated neurotoxicity is dependent on a phylogenetically conserved Asp, whereas the capacity to suppress non-fibrillar, amorphous protein aggregation is not affected by mutating this Asp to Asn. Moreover, conformational changes occur as a result of Asp to Asn mutations in both proSP-C and Bri2 BRICHOS, those changes can partly be mimicked by lowered pH and the conserved Asp titrates with an apparent p*K*a of about 6.5 in both BRICHOS domains studied.

The Asp residue studied herein is the only conserved non-Cys residue in all known BRICHOS domains and two mutations of this residue in human proSP-C BRICHOS (D105) are linked to ILD(16, 20, 36). Based on molecular dynamic simulation(16), monomeric wildtype proSP-C BRICHOS and the D105N mutant behaved differently: only minor conformational changes were seen in the mutant, but several large-scale changes occurred in the wildtype at moderately elevated temperatures, which resulted in a more loosely folded structure. We can rationalize our results against this background (Fig. 6a **and** b). D105N mutation in rh proSP-C BRICHOS indeed results in a more ordered conformation and apparently more efficient trimer formation, while D148N mutation of rh Bri2 BRICHOS results in more readily transition of monomers to more compact dimers (Supplementary Fig. 5a **and** f). Similar effects as observed in the mutants were seen for the wildtype proteins when pH was lowered to 6-7 (Fig. 2h **and** i). This suggests that a negatively charged Asp side-chain is necessary for maintaining a “loose” flexible state of the BRICHOS subunit and that protonation, or replacement with a neutral Asn, results in a more “compact” state that is prone to form oligomers. In the open conformation, BRICHOS is efficient in alleviating Aβ42 amyloid neurotoxicity, while the compact conformation is more potent against overall amyloid fibril formation but inefficient against amyloid induced neurotoxicity (Fig. 6b). Interestingly, in brains of AD patients the pH is lower than that in the brains of healthy individuals(53), and at low pH Aβ can form fibrils more efficiently than at neutral pH(54).

**Fig. 6.**
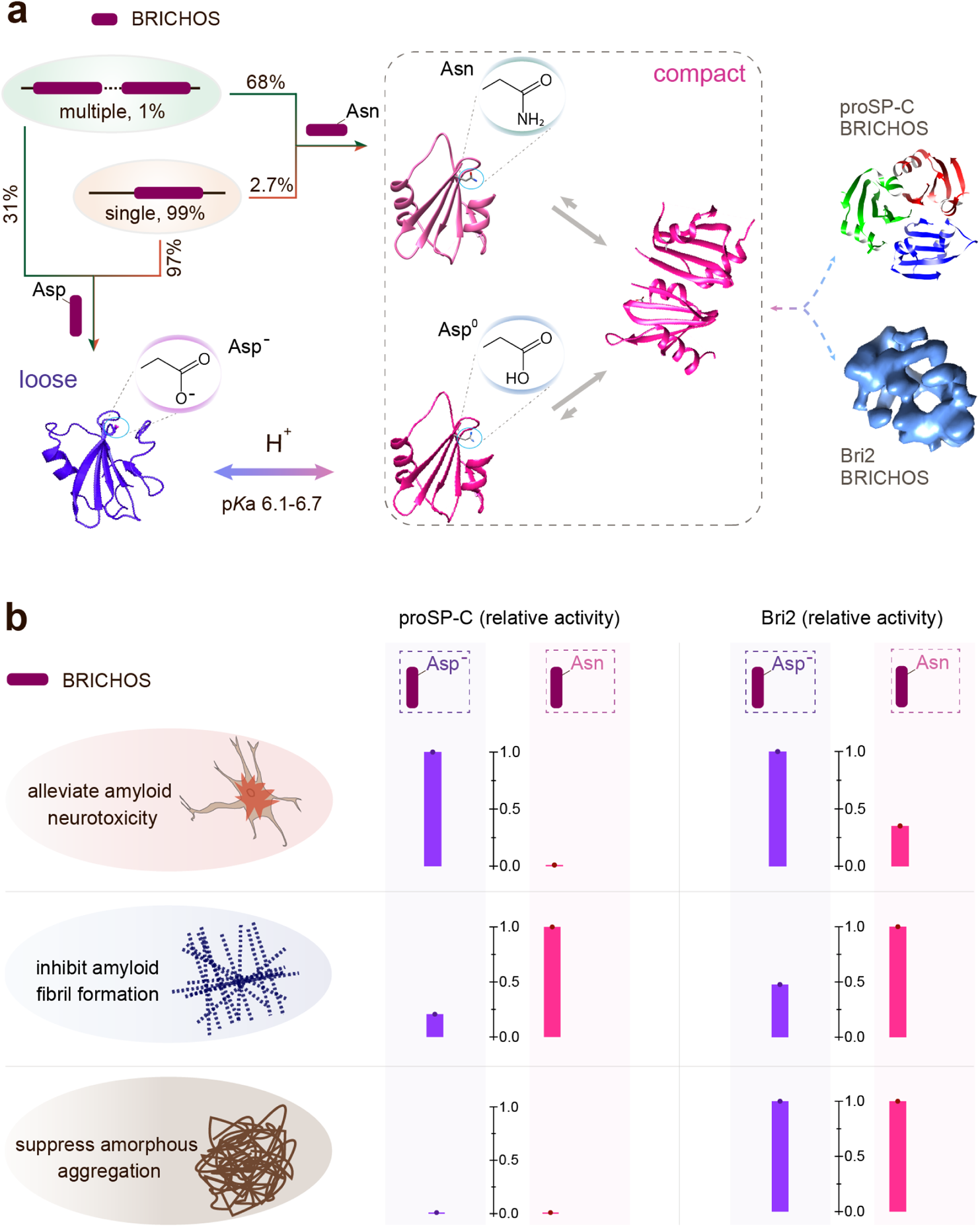
Schematic summary of BRICHOS activities depending on ionization state of its conserved Asp. **(a)** Asp and Asn in BRICHOS domains (single and multiple BRICHOS precursors account for 94% and 5%, and adopt “loose” and “compact” conformation, respectively. At neutral pH, the conserved Asp is ionized (Asp^−^) and BRICHOS presents an open structure (from ref^(16)^). At pH 6-7, the Asp is protonated and BRICHOS is more compact (PDB accession number 2YAD, from crystals prepared at pH 6.5^(ref(16)^) and prone to dimerize, which could be mimicked by Asn present in some BRICHOS families. Eventually, proSP-C BRICHOS forms trimers (2YAD), while Bri2 BRICHOS forms larger oligomeric assemblies (EMD-3918). **(b)** The activities of BRICHOS are regulated by the conserved Asp; with its ionization form BRICHOS can efficient present anti-amyloid-induced neurotoxicity. Protonation of the Asp or replaced by Asn diminishes the activity for BRICHOS anti-amyloid-induced neurotoxicity, but oppositely, the capacity inhibiting amyloid fibril is significantly enhanced. After forming large oligomers, the general chaperone activity against amorphous aggregation is not obviously affected. The y-axis is the normalized activity with substrate alone as 0 and the higher activity from the wildtype species or the mutant as 1. For the neurotoxicity, the data used are from a ratio of 1:2 (substrate: BRICHOS) for proSP-C BRICHOS and 1:1 for Bri2 BRICHOS. For the activity against amyloid fibril formation, the data are from a ratio of 1:1 (substrate: BRICHOS) for proSP-C BRICHOS and 1:0.5 for Bri2 BRICHOS. For the general chaperone activity against amorphous aggregation, the data used are from a ratio of 1:2.4 (substrate: BRICHOS), both for proSP-C BRICHOS and for Bri2 BRICHOS.

The crystal structure of rh proSP-C BRICHOS, the only available high-resolution structure of a BRICHOS domain, gives limited information about the local environment of the conserved Asp since electron density is lacking for the part N-terminally of the juxtaposed helix 2(16). However, an inspection of amino acid sequence alignments of BRICHOS domains from all families show that there is a well conserved carboxylate residue situated just upstream of helix 2 (Supplementary Fig. 3, positions 176-180), and its location close to the sidechain of the strictly conserved Asp may contribute to the apparently elevated p*K*a values of D105 and D148 now observed. The distant evolutionary relationship between Bri2 and proSP-C BRICHOS domains, with <20% sequence identities, makes it likely that the common effects now observed between them also apply to other BRICHOS domains that exhibit similar evolutionary distances. Therefore, an elevated p*K*a value of the only strictly conserved non-Cys residue, as now found for proSP-C and Bri2 BRICHOS, is probably relevant for a common function of all BRICHOS domains, which thus likely requires a maintained “loose” conformation. Our results indicate that such a hypothetical common function is likely to be related to prevention of amyloid toxicity rather than prevention of amorphous protein aggregation. So far, molecular chaperones or chaperone-like domains have not been much studied extensively as regards sensitivity to pH. One exception is clusterin, whose activity is enhanced at mildly acidic pH, which appears to result from an increase in regions with solvent-exposed hydrophobicity, but independent of any major changes in secondary or tertiary structure(55). The phylogenetic analyses showed the presence of the BRICHOS domain in a broad array of proproteins and species. In some cases where multiple BRICHOS domains are found, the conserved Asp is preferentially replaced with Asn (Fig. 1 **and** 2). Considering the experimental results herein this may suggest that BRICHOS has adopted different types of molecular chaperone activities during evolution.

Molecule chaperones are essential guards of all living cells, while the chaperoning abnormality may cause disease—chaperonopathy. The chaperoning capacities of BRICHOS domain against amyloid neurotoxicity and fibril formation can apparently be modulated by a conserved Asp in response to pH changes. In the human brain, pH decreases with aging(56), and also pH is significantly lower in patients than in heathy controls in different human brain disorders(53, 57). The observed phenomenon in this study suggests the possibility that microenvironmental changes may lead to human disease. The results here are based on *in vitro* and *ex vivo* experiments, and further *in vivo* work that focuses on the activities of BRICHOS and pH-mediated regulation can shed further light on this.

## Materials and Methods

### Phylogenetic analysis of BRICHOS domain

From the SMART database(58), 3 355 BRICHOS sequences were downloaded while the sole bacterial BRICHOS precursor (*Paeniclostridium sordellii* ATCC 9714) was not included. Incomplete sequences were filtered out, resulting in total of 3 190 sequences. Identical sequences were filtered by means of CD-HIT(59), with a threshold of 100% of sequence identity, which gave 3 093 amino acid sequences. The hidden markov model profile (HMM) of BRICHOS (PF04089) was extracted from PFAM database(60). The CD-HIT filtered BRICHOS protein sequences were then scanned against the HMM profile using HMMER software v3.3.2(61) with an E-value cut-off less than 1.0×10^-5^. An in-house python script was written to filter out the significant BRICHOS proteins from the HMM result file and to extract the BRICHOS domain sequences (each BRICHOS amino acid sequence was extended six residues upstream from the BRICHOS domain starting position defined by PFAM). Further, BRICHOS domain sequences less than 68 aa were removed as one rodent BRICHOS domain is just 69 residues and still functional against amyloid fibril formation(62), which eventually generated 2 019 BRICHOS sequences. The multiple sequence alignment of the 2019 BRICHOS domain sequences were created using MAFFT alignment server with the default settings(63), and the sequence logo was generated using Weblogo webserver(64). RAxML HPC(v8.2.10)(65) was employed for constructing the phylogenetic tree using the PROTGAMMAAUTO model with 100 times bootstrap iterations. The tree shown in this study was visualized using the Interactive Tree of Life (iTOL) server(66) and Geneious software. The taxonomy tree common for species that contain BRICHOS precursors were generated by NCBI Taxonomy (https://www.ncbi.nlm.nih.gov/Taxonomy/CommonTree/wwwcmt.cgi) and visualized by Geneious software.

### Rh Bri2 BRICHOS and rh proSP-C BRICHOS wildtype and mutant preparation

For generating rh Bri2 BRICHOS D148N, the amplification primers 5’- ATAGTGATCCTGCCAACATTGATAACTTTAACAAGAAACTTACA-3’ and 5’- TGTAAGTTTCTTGTTAAAGTTATGAACAATGTTGGCAGGATCACTAT-3’ were synthesized. With the wildtype NT*-Bri2 BRICHOS (corresponding to the solubility tag NT* followed by Bri2 residues 113–231(32, 67)) plasmid as PCR template Bri2 BRICHOS D148N was obtained with QuikChange II XL Site-Directed Mutagenesis Kit (Agilent, US), and the DNA sequence was confirmed (GATC Bioteq, Germany). Similarly, the amino acid Thr at position 206 was mutated to Trp using KAPA HiFi HotStart ReadyMix PCR Kit (Kapa Biosystems, USA) together with the designed complementary primers (5’- CCTATCTGATTCATGAGCACATGGTTATTTGGGATCGCATTGAAAAC-3’ and 5’- GTTTTCAATGCGATCCCAAATAACCATGTGCTCATGAATCAGATAGG-3’) and verified by sequencing (Eurofins Genomics). As described(32, 33) , the Bri2 BRICHOS variants were expressed in SHuffle T7 *E. coli* cells. Briefly, the cells were incubated at 30°C in LB medium with 15 µg mL^-1^ kanamycin until an OD_600 nm_ around 0.9. For overnight expression, the temperature was lowered to 20°C, and 0.5 mmol L^-1^ (final concentration) isopropyl β-D-1-thiogalactopyranoside (IPTG) was added. The induced cells were harvested by centrifugation (4°C, 7 000 ×g) and the cell pellets were resuspended in 20 mmol L^-1^ Tris pH 8.0. After 5 min sonication (2 s on, 2 s off, 65% power) on ice, the lysate was centrifuged (4°C, 24 000 ×g) for 30 min and the protein of interest was isolated with a Ni-NTA column. To remove the His_6_-NT* part, the target proteins were cleaved with thrombin (1:1 000, w/w) at 4°C overnight and loaded over a second Ni-NTA column. Different species of rh Bri2 BRICHOS variants were isolated and analysed by Superdex 200 PG, 200 GL or 75 PG columns (GE Healthcare, UK) using an ÄKTA system (GE Healthcare, UK) with buffer of 20 mmol L^-1^ NaPi (Sodium Phosphate) with 0.2 mmol L^-1^ EDTA at different pHs. For generating rh proSP-C BRICHOS D105N, the PCR primers 5’- CACTGGCCTCGTGGTGTATAACTACCAGCAGCTGCTGATCGC-3’ and 5’- GCGATCAGCAGCTGCTGGTAGTTATACACCACGAGGCCAGTG -3’ were synthesized. With the wildtype proSP-C BRICHOS (corresponding to the solubility tag thioredoxin followed by proSP-C residues 86–197(16)) plasmid as PCR template, proSP-C BRICHOS D105N was obtained with QuikChange II XL Site-Directed Mutagenesis Kit (Agilent, US), and the DNA sequence was confirmed (GATC Bioteq, Germany). The expression and purification were performed as described(16, 68). Briefly, both wildtype proSP-C BRICHOS and the D105N mutant were expressed in Origami 2 (DE3) pLysS *E. coli* cells. The cells were grown at 37°C in LB medium containing 100 μg mL^-1^ ampicillin until an OD_600 nm_ around 0.9. The temperature was lowered to 25°C and 0.5 mmol L^-1^ (final concentration) IPTG was added for overnight expression. The cells were harvested by centrifugation at 7 000 × *g* for 20 min, and the cell pellets were resuspended in 20 mmol L^-1^ Tris pH 8.0. The protein was purified using Ni-NTA column and ion exchange chromatography (QFF, GE Healthcare). Thrombin (1:1 000, w/w) was used to remove the thioredoxin tag. The purified rh proSP-C BRICHOS variants were analysed by Superdex 200 GL columns (GE Healthcare, UK) using an ÄKTA system (GE Healthcare, UK). The BRICHOS mutants in this study were expressed and purified in parallel with their wildtype counterparts.

### NMR spectroscopy

For the NMR experiments, gene fragment encoding human proSP-C BRICHOS was transformed into SHuffle T7 *E. coli* cells and was grown in LB with gradually increasing D_2_O content (25%, 50%,75% and 100%). At 100% D_2_O, 1 mL LB was used to inoculate 100 mL M9 in 100% D_2_O enriched with ^15^N H_4_Cl and ^13^C glucose, and was grown over night at 31°C. After overnight incubation, the 100 mL was added to 900 mL M9 in D_2_O enriched with ^15^N and ^13^C and grown at 30°C until OD_600 nm_ was around 0.8. The temperature was lowered to 20°C and 0.5 mmol L^-1^ (final concentration) IPTG was added for overnight expression. The purification was performed as described above.

2D ^1^H-^15^N TROSY-HSQC experiments were obtained at 37°C on Bruker 800 MHz spectrometer equipped with a TCI cryogenic probe. Spectra were processed with the software NMRPipe and visualized using Sparky NMR. The concentrations of ^2^H, ^15^N,^13^C-labeled proSP-C was 288 μmol L^-1^ in 20 mmol L^-1^ ammonium acetate pH 7.2 and 229 μmol L^-1^ in 20 mmol L^-1^ ammonium acetate pH 5.5, both in 90% H_2_O/10% D_2_O.

### Circular dichroism and fluorescence spectroscopy and aggregation analyses

CD spectra were recorded in 1 mm path length quartz cuvettes at 25°C from 260 to 185 nm in a J-1500 Circular Dichroism Spectrophotometer (JASCO, Japan) with a protein concentration of 4.2–10 µmol L^-1^. The bandwidth was set to 1 nm, data pitch 0.5 nm, and scanning speed 50 nm min^-1^. The spectra shown are averages of five consecutive scans.

Citrate synthase from porcine heart (Sigma-Aldrich, Germany) was diluted in 40 mmol L^-1^ HEPES/KOH pH 7.5 to 600 nmol L^-1^ (calculated from a molecular weight of 85 kDa corresponding to a dimer(69)) and then equilibrated at 45°C with and without different concentrations of rh Bri2 BRICHOS D148N oligomer or proSP-C BRICHOS variants. The aggregation kinetics were measured by reading the apparent increase in absorbance at 360 nm under quiescent conditions using a microplate reader (FLUOStar Galaxy from BMG Labtech, Offenberg, Germany).

One µmol L^-1^, calculated for the monomeric subunit, of different rh BRICHOS in 20 mmol L^-1^ NaPi pH 8.0 or pH 6.0 were incubated at 25°C with 2 µmol L^-1^ bis-ANS (4,4’-Bis(phenylamino)-[1,1’-binaphthalene]-5,5’-disulfonic acid dipotassium salt) for 10 min, and the fluorescence emission spectra were recorded from 420 to 600 nm after excitation at 395 nm with the Infinite M1000 plate reader (Tecan, Austria). Rh Bri2 BRICHOS T206W monomers were diluted to 2 μmol L^-1^ by using 20 mmol L^-1^ NaPi containing 0.2 mmol L^-1^ EDTA with different pH in the final samples in the range of 6.3–8.0. For tryptophan fluorescence measurements, samples were prepared in duplicates with a volume of 150 μL. Samples were excited at 280 nm (5 μm bandwidth) and fluorescence emission from 300–400 nm (10 μm bandwidth, 1 nm step interval) was recorded on black polystyrene flat-bottom 96-well plates (Costar) using a spectrofluorometer (Tecan Saphire 2). For the final fluorescence intensities, the results were corrected by subtracting the background fluorescence of the buffer.

### Transmission electron microscopy imaging of rh Bri2 BRICHOS D148N oligomers and single particle processing

Rh Bri2 BRICHOS D148N oligomers after SEC isolation were immediately stored on ice followed by grid preparation. Aliquots (4 µL) were adsorbed onto glow-discharged continuous carbon-coated copper grids (400 mesh, Analytical Standards) for one min. The grids were subsequently blotted with filter paper, washed with two drops of milli-Q water, and negatively stained with one drop of 2% (w/v) uranyl acetate for 45 s before final blotting and air-drying. The sample was imaged using a Jeol JEM2100F field emission gun transmission electron microscope (Jeol, Japan) operating at 200 kV. Single micrographs for evaluating the quality of the sample were recorded on a Tietz 4k × 4k CCD camera, TVIPS (Tietz Video and Image Processing Systems, GmbH, Gauting, Germany) at magnification of× 85 200 (1.76Å per pixel) and 1.0–2.8 µm defocus. A total of 16 micrographs were recorded for single particle analysis. All 16 micrographs were imported to EMAN2 (version 2.3) for further processing(70). After importing and estimating defocus with e2evalimage.py, single particles in different orientations were selected from the images using e2boxer.py in manual mode (11 094 particles, after one more manual selection, 10 223 particles were left). For each image, the contrast transfer function (CTF) parameters were estimated on boxed out regions (containing particles, 168×168) using e2ctf.auto.py program. A reference-free 2D classification based on the selected 10 223 phase-flipped particles (low-pass filtered to 20 Å) was performed using e2refine2d.py. The 2D classes show an approximate 2-fold symmetry, which is consistent with both the results of rh wildtype Bri2 BRICHOS oligomer and the biochemical data. Generated 2D classes were used as the input for building the 3D initial model using e2initialmodel.py. 3D refinement was performed in several rounds using e2refine_easy.py applying D2 symmetry aiming at a final resolution of 15 Å. The first two rounds of 3D refinements were performed with pixel size of 3.52 Å after binning the data by a linear factor 2. In the last round of refinement, the data was resampled to 2.464 Å per pixel. The final map from the first round of refinements was used as model in the second, and the final map from the second round of refinements was used as model in the third. The resolution was determined based on a Fourier shell correlation (FSC) value of 0.143(71), following the gold standard FSC procedure implemented in EMAN2(72).

### BRICHOS and Aβ42 monomer interaction monitored by surface plasmon resonance

Aβ42 monomers were immobilized by amine coupling onto flowcell 4 on a CM5 sensor chip (GE Healthcare) using a BIAcore 3000 instrument (BIAcore AB). A reference surface was prepared on flowcell 3 using the same coupling protocol but without protein injected. The immobilization was performed at a flow rate of 20 μL min^-1^ with 20 mmol L^-1^ sodium phosphate pH 8.0 containing 0.2 mmol L^-1^ EDTA as running buffer, and the other details were set according to the manufacturer’s instructions. After immobilization with a final reponse level of 727 RU, the flow-cells were stabilized over night in running buffer (10 mmol L^-1^ HEPES pH 7.5 containing 150 mmol L^-1^ NaCl and 0.2 mmol L^-1^ EDTA) at a flow rate of 20 μL min^-1^. For binding analysis, 25 μmol L^-1^ rh wildtype proSP-C BRICHOS or the D105N mutant in 10 mmol L^-1^ HEPES pH 7.5 containing 150 mmol L^-1^ NaCl and 0.2 mmol L^-1^ EDTA were injected in over the chip surfaces at 25°C at a flow rate of 20 μL min^-1^ for 3 min, respectively. 10 mmol L^-1^ NaOH was used for further chip surface regeneration. For kinetic analysis, different concentrations of rh proSP-C BRICHOS D105N mutant in running buffer, *i.e.* 0, 1.56, 3.13, 6.25, 12.5, 25, 50 and 100 μmol L^-1^, were individually injected in over the chip surfaces at 25°C at a flow rate of 20 μL min^-1^. 30 mmol L^-1^ NaOH was used for further chip surface regeneration. In all experiments, the response from the blank surface was subtracted from the immobilized surface response, the baselines of the sensorgrams were adjusted to zero and buffer spikes were excluded.

Steady state apparent affinities for rh proSP-C BRICHOS D105N mutant to immobilised Aβ42 monomers were estimated by plotting the maximum binding response versus BRICHOS concentrations. The baseline of the sensorgrams were adjusted to zero and buffer spikes were excluded for global fits to reflect the binding affinity. Since the response signals of the two lowest protein concentrations (*i.e.,* 1.56 and 3.13 μmol L^-1^) used in kinetic analysis were too weak, only sensorgrams obtained from rh proSP-C BRICHOS D105N mutant ranging from 6.25 μmol L^-1^ to 100 μmol L^-1^ were included in the global fits. The dissociation was globally fitted to a biexponential model as described by Eq. (**1**)(28, 73):

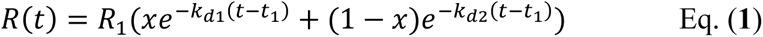

where *R_1_* is the response signal at the starting time for the dissociation phase *t*_1_, and *x* is between 0 and 1. *k_d1_* and *k_d2_* are the dissociation rate constants for the fast and slow dissociation phases, respectively. The global fitted value *k_d_*_2_ was used to calculate the apparent *K*_D_ value.

The association phase was fitted to Eq. (**2**)(28):

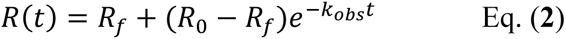

where *R_0_* and *R_f_* are the initial and final response signal of the association phase, respectively. *k_obs_* is the observed rate constant described by Eq. (**3**)(28, 73):

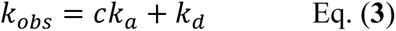

where *c* is the protein concentration, *k_a_* is the association rate constant, and *k_d_* from this analysis is related to secondary binding artifacts corresponding to *k_d_*_1_. Linear regression was used to determine *k_a_*. The apparent *K*_D_ value was calculated as ratio of the dissociation rate constant and association rate constant.

### Aβ42 monomer preparation and ThT assay

Recombinant Met-Aβ(1–42), here referred to as Aβ42, was produced in BL21*(DE3) pLysS *E. coli* (B strain) cells and purified by ion exchange(32). The purified Aβ42 peptides were lyophilized and re-dissolved in 20 mmol L^-1^ Tris pH 8.0 with 7 mol L^-1^ Gdn-HCl, and the monomers were isolated in 20 mmol L^-1^ sodium phosphate pH 8.0 with 0.2 mmol L^-1^ EDTA by a Superdex 30 column 26/600 (GE Healthcare, UK). The concentration of monomeric Aβ42 was calculated with using an extinction coefficient of 1 424 M^-1^ cm^-1^ for (A_280_−A_300_). For Aβ42 fibrillization kinetics analysis, 20 µL solution containing 10 µmol L^-1^ ThT, 3 µmol L^-1^ Aβ42 monomer and different concentrations of rh BRICHOS at molar ratios 0, 10, 30, 50, 70 or 100 % (relative to Aβ42 monomer molar concentration), were added to each well of half-area 384-well microplates with clear bottom (Corning Glass 3766, USA), and incubated at 37°C under quiescent conditions. The ThT fluorescence was recorded using a 440 nm excitation filter and a 480 nm emission filter using a microplate reader (FLUOStar Galaxy from BMG Labtech, Offenberg, Germany). For Aβ42 seeds preparation, 3 µmol L^-1^ Aβ42 monomers were incubated for about 20 h at 37°C, and the fibrils were then sonicated in a water bath for 3 min. For seeding of Aβ42 fibrillization, 20 µL solution containing 10 µmol L^-1^ ThT, 3 µmol L^-1^ Aβ42, different concentrations of rh BRICHOS at 0, 10, 50, 70 and 100 %, and 0.6 µmol L^-1^ seeds (calculated from the original Aβ42 monomer concentration) were added at 4°C to each well in triplicate of 384-well microplates with clear bottom (Corning Glass 3766, USA) and immediately incubated at 37°C under quiescent conditions. The elongation rate constant *k_+_* in the presence of rh BRICHOS was calculated from the highly seeded experiments. The initial slope of the concave aggregation traces was determined by a linear fit of the first ∼20 min traces. For all the experiments, aggregation traces were normalized and averaged using four or three replicates, and data defining one dataset was recorded from the same plate at the same time, and all the ThT data was from the same batch of Aβ42 peptide.

### Analysis of Aβ42 aggregation kinetics

Fibrillization traces of Aβ42 with and without different concentrations of rh BRICHOS were fitted to a sigmoidal equation Eq. (**4**)(32, 33), where the half time *τ*_1/2_ and the maximal growth rate *r_max_* were extracted:

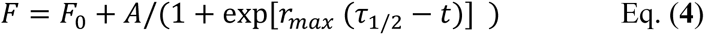

where *A* the amplitude and *F_0_* the base value.

To dissect the molecular mechanism underlying BRICHOS counteracting Aβ42 aggregation, the fibrillization traces were globally fitted by Eq. (**5**)(27):

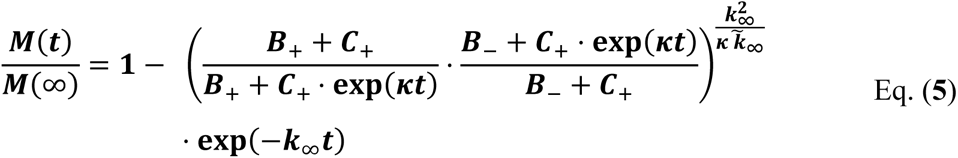

where *M(t)* is the total fibril mass concentration, and the intermediate coefficients are functions of λ and κ, and *n_C_* and *n_2_* are the reaction orders for primary and secondary nucleation, respectively:

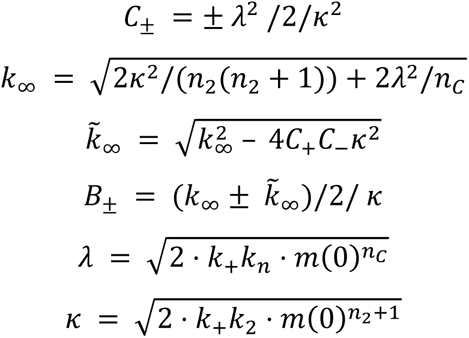

The microscopic rate constants *k_n_*, *k_+_*, and *k_2_* are for primary nucleation, elongation, and secondary nucleation, respectively. The kinetic data were globally fitted to Eq. (**5**), where the fits were partially constrained with one fitting parameter held to a constant value, resulting in that only one rate constant (*k_n_*, *k_+_* or *k_2_*) is the sole fitting parameter(32, 33). To investigate the generation of nucleation units, according to the nucleation rate *r_n_(t)*(27):

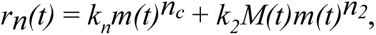

the total number of nucleation units was calculated by integrating the nucleation rate *r_n_(t)* over the reaction.

### Immunogold staining of Aβ42 fibrils and transmission electron microscopy analysis

Five µmol L^-1^ Aβ42 monomer was incubated at 37°C with 100% rh wildtype proSP-C BRICHOS and the D105N mutant overnight, and the fibrils were collected at 4°C by centrifugation for 1 h at 22 000×g. The fibrils were gently resuspended in 20 µL 1×TBS, of which 2 µL were applied to carbon coated copper grids, and incubated for about 5 min. Excess solution was removed and the girds were blocked by incubation in 1% BSA in 1×TBS for 30 min, followed by 3×10 min washing by 1×TBS. The grids were then incubated with polyclonal antibody against human proSP-C (SFTPC) (1:200 dilution, Atlas Antibodies) overnight at 4°C, and washed 3×10 min with 1×TBS. Finally, the grids were incubated with anti-rabbit IgG-gold coupled to 20 nm gold particles (1:40 dilution, BBI Solutions) for 2 h at room temperature, and washed 5×10 min with 1×TBS. Excess solution was removed, and 2 µL of 2.5% uranyl acetate was added to each grid (kept about 20 s). Excess solution was removed, and the grids were air-dried at room temperature, and analysed by transmission electron microscopy (TEM, Jeol JEM2100F at 200 kV).

### Electrophysiological recordings

All the animal experiments were carried out in accordance with the ethical permit granted by Norra Stockholm’s Djurförsöksetiska Nämnd (dnr N45/13). C57BL/6 mice of either sex (postnatal days 14–23, supplied from Charles River, Germany) were used in the experiments. Before sacrificed, all the mice were anaesthetized deeply using isoflurane.

The brain was dissected out and placed in modified ice-cold ACSF (artificial cerebrospinal fluid). The ACSF contained 80 mmol L^-1^ NaCl, 24 mmol L^-1^ NaHCO_3_, 25 mmol L^-1^ glucose, 1.25 mmol L^-1^ NaH_2_PO_4_, 1 mmol L^-1^ ascorbic acid, 3 mmol L^-1^ NaPyruvate, 2.5 mmol L^-1^ KCl, 4 mmol L^-1^ MgCl_2_, 0.5 mmol L^-1^ CaCl_2_ and 75 mmol L^-1^ sucrose. Horizontal sections (350 µm thick) of the ventral hippocampi from both hemispheres were sliced with a Leica VT1200S vibratome (Microsystems, Sweden). The sections were immediately transferred to a submerged incubation chamber containing standard ACSF: 124 mmol L^-1^ NaCl, 30 mmol L^-1^ NaHCO_3_, 10 mmol L^-1^ glucose, 1.25 mmol L^-1^ NaH_2_PO_4_, 3.5 mmol L^-1^ KCl, 1.5 mmol L^-1^ MgCl_2_ and 1.5 mmol L^-1^ CaCl_2_. The chamber was held at 34°C for at least 20 min after dissection and it was subsequently cooled to room temperature (∼22°C) for a minimum of 40 min. Proteins (Aβ42 and rh BRICHOS) were first added to the incubation solution for 15 min, and then the slices were transferred to the interface-style recording chamber for extracellular recordings. During the incubation, slices were supplied continuously with carbogen gas (5% CO_2_, 95% O_2_) bubbled into the ACSF.

Recordings were performed with borosilicate glass microelectrodes in hippocampal area CA3, pulled to a resistance of 3–5 MΩ, filled with ACSF and placed in stratum pyramidale. Local field potentials (LFP, γ oscillations) were recorded at 32°C in an interface-type chamber (perfusion rate 4.5 mL per minute) and elicited by applying kainic acid (100 nmol L^-1^, Tocris). The oscillations were stabilized for 20 min before any recordings. No Aβ42, rh Bri2 BRICHOS R221E species or combinations thereof were present in the recording chamber either during γ oscillations stabilization, or during electrophysiological recordings. The interface chamber recording solution contained 124 mmol L^-1^ NaCl, 30 mmol L^-1^ NaHCO_3_, 10 mmol L^-1^ glucose, 1.25 mmol L^-1^ NaH_2_PO_4_, 3.5 mmol L^-1^ KCl, 1.5 mmol L^-1^ MgCl_2_ and 1.5 mmol L^-1^ CaCl_2_.

Interface chamber LFP recordings were carried out by a 4-channel amplifier/signal conditioner M102 amplifier (Electronics lab, University of Cologne, Germany). The signals were sampled at 10 kHz, conditioned using a Hum Bug 50 Hz noise eliminator (LFP signals only; Quest Scientific, North Vancouver, BC, Canada), software low-pass filtered at 1 kHz, digitized and stored using a Digidata 1322A and Clampex 9.6 software (Molecular Devices, CA, USA).

Power spectral density plots (from 60 s long LFP recordings) were calculated using Axograph X (Kagi, Berkeley, CA, USA) in averaged Fourier-segments of 8 192 points. Oscillation power was calculated from the integration of the power spectral density from 20 to 80 Hz.

### Statistics and reproducibility

The electrophysiological data is presented as means ± standard errors of the means. Prior statistical analysis all the data was subjected to outlier determination and removal with the ROUT method (Q = 1%) followed by D’Agostino & Pearson omnibus normality test. Based on the previous experience with the outliers and overall sample behaviour, sample size was determined based on previous studies performed in an interface-type chamber(27, 32, 33, 47, 74, 75). Each experimental round was performed with parallel controls (Control KA and Aβ42) from the same animal and randomized preparations (slices incubated with Aβ42 + rh proSP-C BRICHOS D105N, + wildtype rh proSP-C BRICHOS, + rh Bri2-BRICHOS D148N monomers or + wildtype rh Bri2-BRICHOS). For comparison purposes data from Control KA and Aβ42 was pooled from interleaved slices recorded in these conditions. The number of recorded slices per condition (at least from 3–5 mice) are shown in the corresponding figure legend and source data file. Kruskal-Wallis test followed by Dunn’s multiple comparisons were carried out due to the non-parametric nature- or relatively small size of some data. Comparison of the preventative efficacies of rh Bri2-BRICHOS D148N monomers and rh proSP-C BRICHOS D105N was assessed with the Mann Whitney test. Significance levels are * *p*<0.05, ** *p*<0.01, and \*\*\**p*<0.001. The ThT assay data are presented as means ± standard deviation, and the aggregation traces are averaged from four or three replicates.

## Funding

Olle Engkvists Stiftelse 192-522

Petrus and Augusta Hedlunds Stiftelse M-2018-0998

Swedish Alzheimer Foundation AF-836251

Åhlén-stiftelsens mC9h18

Karolinska Institutet Research Foundation Grant 2020-01819

Stiftelsen för Gamla Tjänarinnor 2019-00779

Loo and Hans Osterman Foundation 2019-01130

Geriatric Diseases Foundation at Karolinska Institutet 2020-02290

Magnus Bergvall Foundation 2019-03010

Swedish Research Council 2020-02434

Swedish Brain Foundation FO2018-0312

Center for Innovative Medicine (CIMED)

Swedish Society for Medical Research P17-0047

FORMAS 2020-01013

Åke Wiberg Foundation M20-0148

JPco-fuND/EU PETABC 2020-02905/EC 643417

FLPP lzp-2018/1-0275

Stiftelsen Sigurd och Elsa Goljes Minne LA2020-0031

## Author contributions

G.C., Y.A.T., X.Z., H.P., H.B., A.L., and N.K. performed experiments. S.H. and G.C. carried out bioinformatic analyses and visualization. G.C., Y.A.T., X.Z., H.B., H.P., A.A., N.K., A.R., H.H., P. K., A.F., and J.J. analysed the data. G.C. and J.J. conceived the study. G.C. and J.J. wrote the paper. All authors discussed the results and commented on the manuscript.

## Competing interests

The authors declare no competing financial interests.

## Data and materials availability

The density map of the Bri2 BRICHOS D148N oligomer have been deposited in the Electron Microscopy Data Bank (EMDB) under the accession code EMD-13005. All data and materials related to this paper are available from G.C.

## Supplementary Figures

**Fig. S1.**
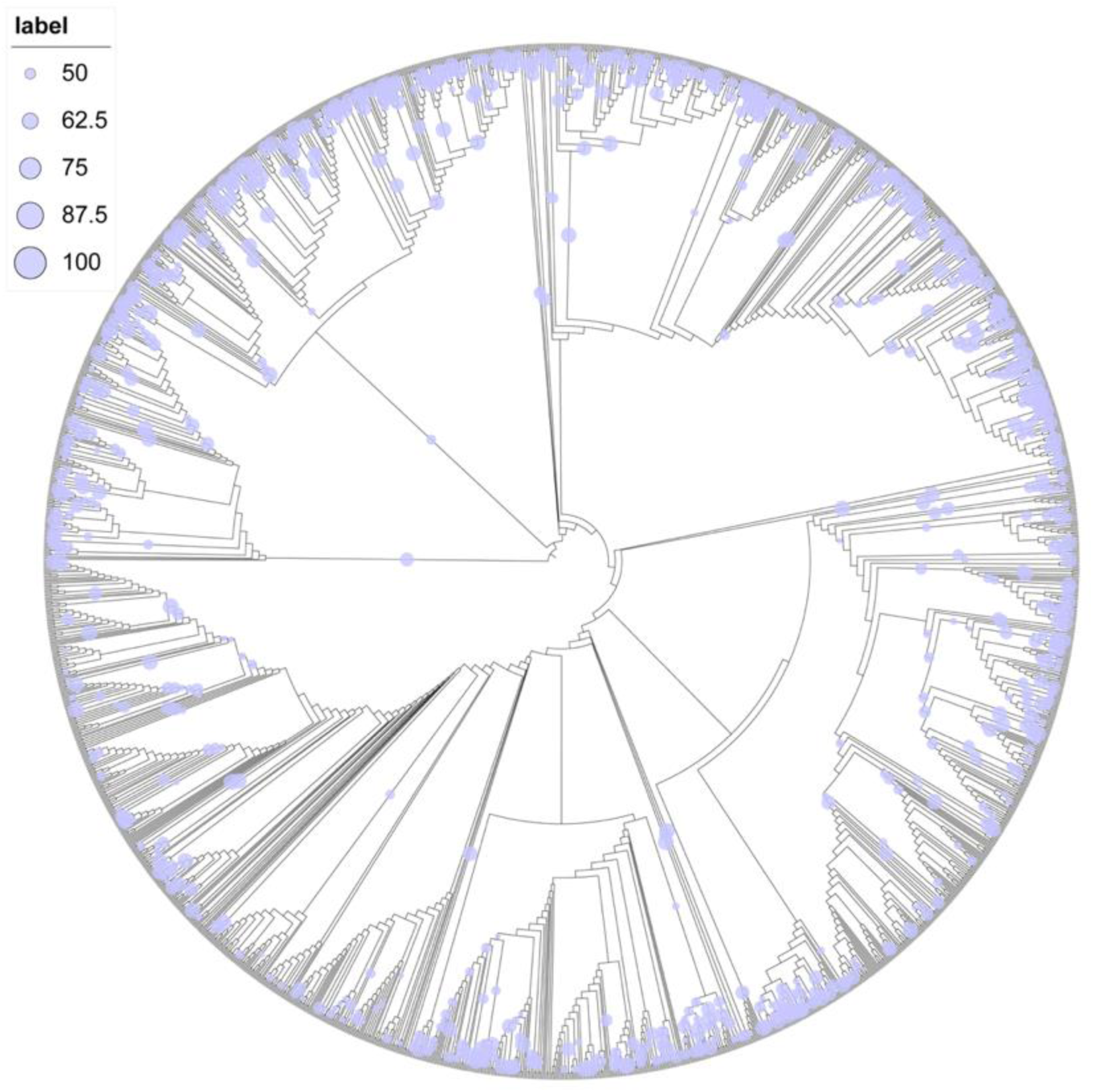
Evolution of the molecular chaperone domain BRICHOS. Phylogenetic tree of the 2 019 BRICHOS domains, which was grouped into thirteen families (Fig. 1c). The bootstraps larger than 50% are indicated by labels, where the sizes are proportional to the bootstrap values.

**Fig. S2.**
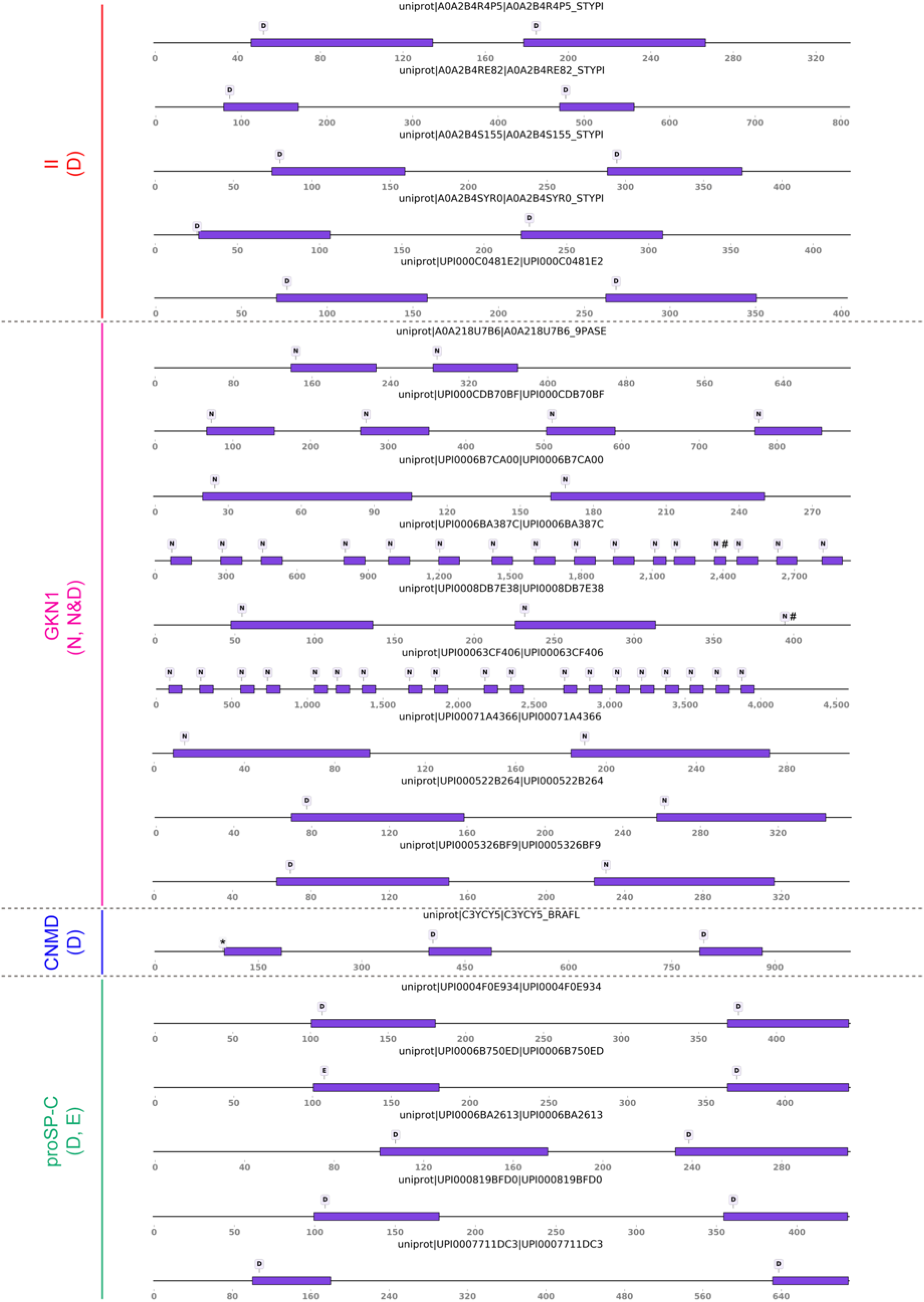
The architecture of proproteins containing multiple BRICHOS domains. The proproteins contain up to nineteen multiple, tandem BRICHOS domains (multi-BRICHOS). The architecture is generally shown in Figure 1d. The uniport accession number is shown over each corresponding cartoon. The length of the bar is not proportional to the size of the BRICHOS domain from different precursors. # refers to the C-terminally truncated, and * indicated the N-terminally truncated.

**Fig. S3.**
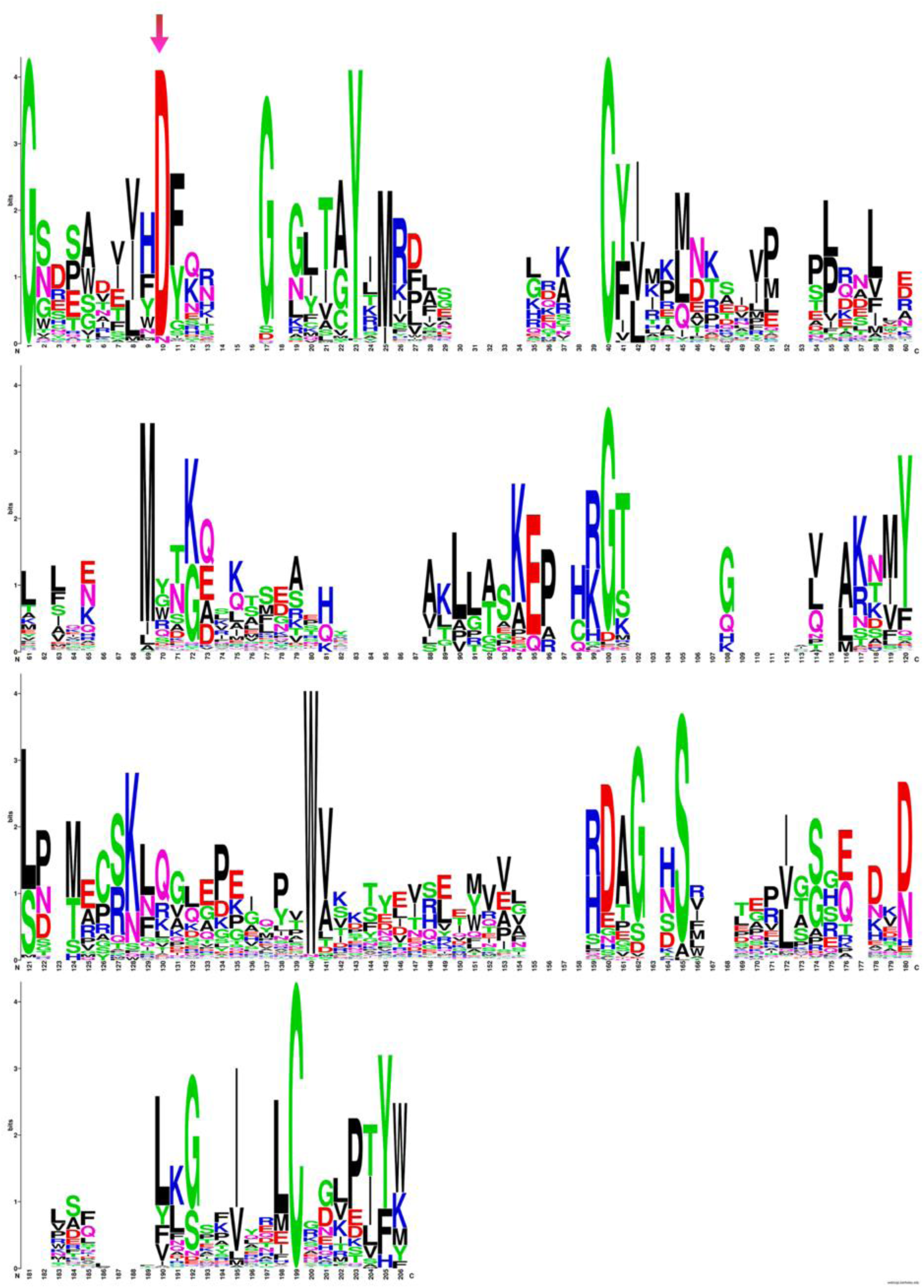
WebLogo representation of the 1 908 single BRICHOS domains. The height of the residue stack implies the sequence conservation, while the height of symbols within the stack indicates the relative frequency of each residue. The arrow marks the conserved Asp residue.

**Fig. S4.**
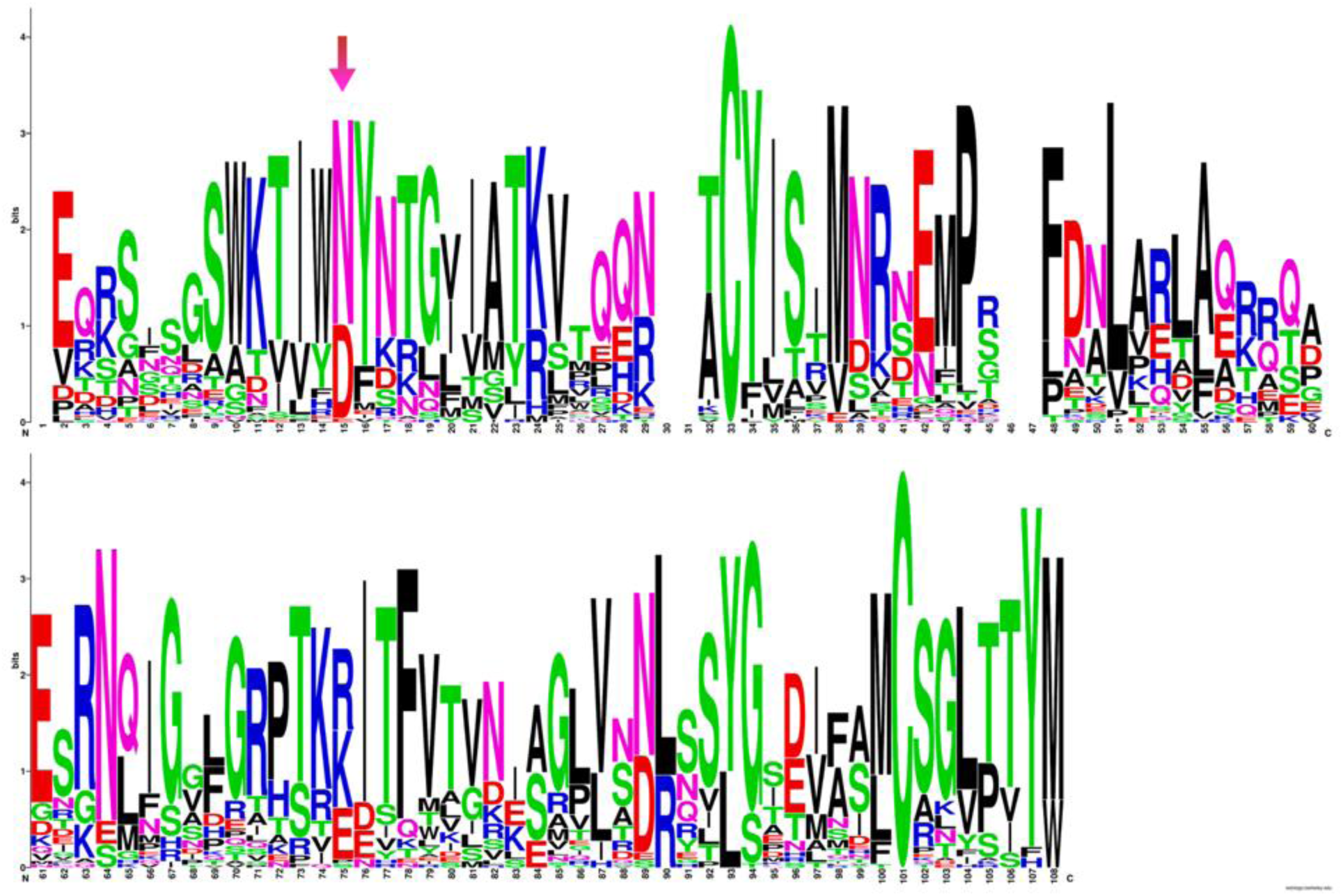
WebLogo representation of the 74 multiple BRICHOS domains. The height of the residue stack implies the sequence conservation, while the height of symbols within the stack indicates the relative frequency of each residue. The arrow marks the conserved Asp residue.

**Fig. S5.**
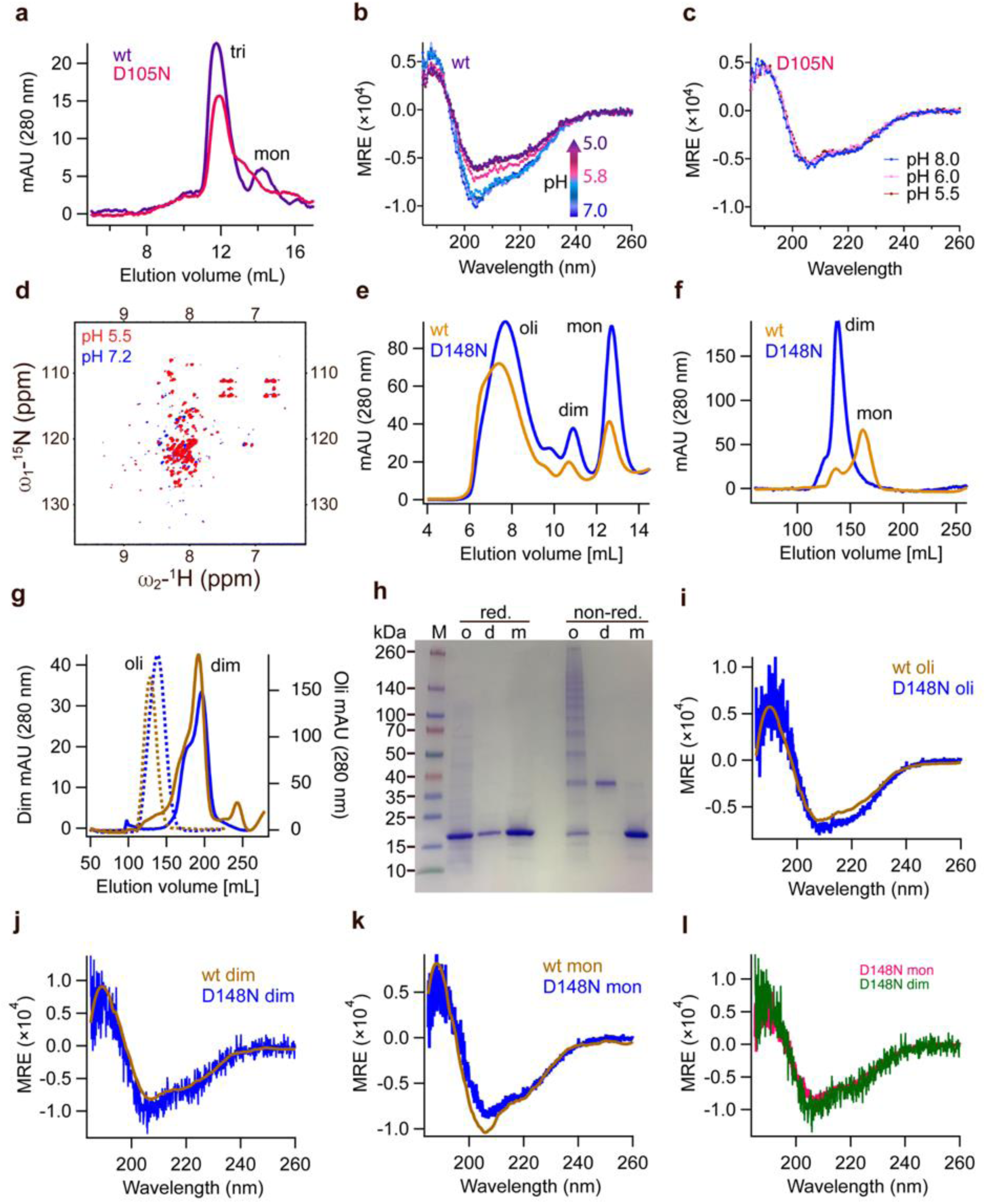
Characterization of rh wildtype and Asp to Asn mutant BRICHOS. (**a**) SEC analysis of purified rh wildtype (wt) proSP-C BRICHOS (purple) and the D105N mutant (red). (**b** and **c**) CD measurements of rh wt proSP-C BRICHOS (purple) and the D105N mutant at different pHs. (**d**) NMR HSQC spectrum comparison of rh wt proSP-C BRICHOS (purple) and the D105N mutant at pH 7.2 (blue) and 5.5 (red). (**e**) SEC of rh wildtype (wt, yellow) NT*-Bri2 BRICHOS and the D148N mutant (blue) species. oli, oligomers; dim, dimers. (**f**) SEC of wildtype (wt) Bri2 BRICHOS (blue) and Bri2 BRICHOS D148N (yellow) monomer fraction prepared from corresponding fusion protein (NT*-Bri2 BRICHOS) monomers. dim, dimers; mon, monomers. (**g**) SEC of wildtype (wt) Bri2 BRICHOS (blue) and the D148N (yellow) oligomer (dash line) and dimer (solid line) fractions prepared from corresponding fusion protein (NT*-Bri2 BRICHOS) oligomer and dimer. oli, oligomers; dim, dimers. (**h**) SDS-PAGE analyses of SEC isolated rh Bri2 BRICHOS D148N species under reducing and non-reducing conditions. (**i–l**) CD spectra of rh Bri2 BRICHOS D148N monomers, dimers and oligomers and the comparison with rh wildtype (wt) Bri2 BRICHOS monomer. MRE is the mean molar residual ellipticity in deg·cm^2^·dmol^-1^.

**Fig. S6.**
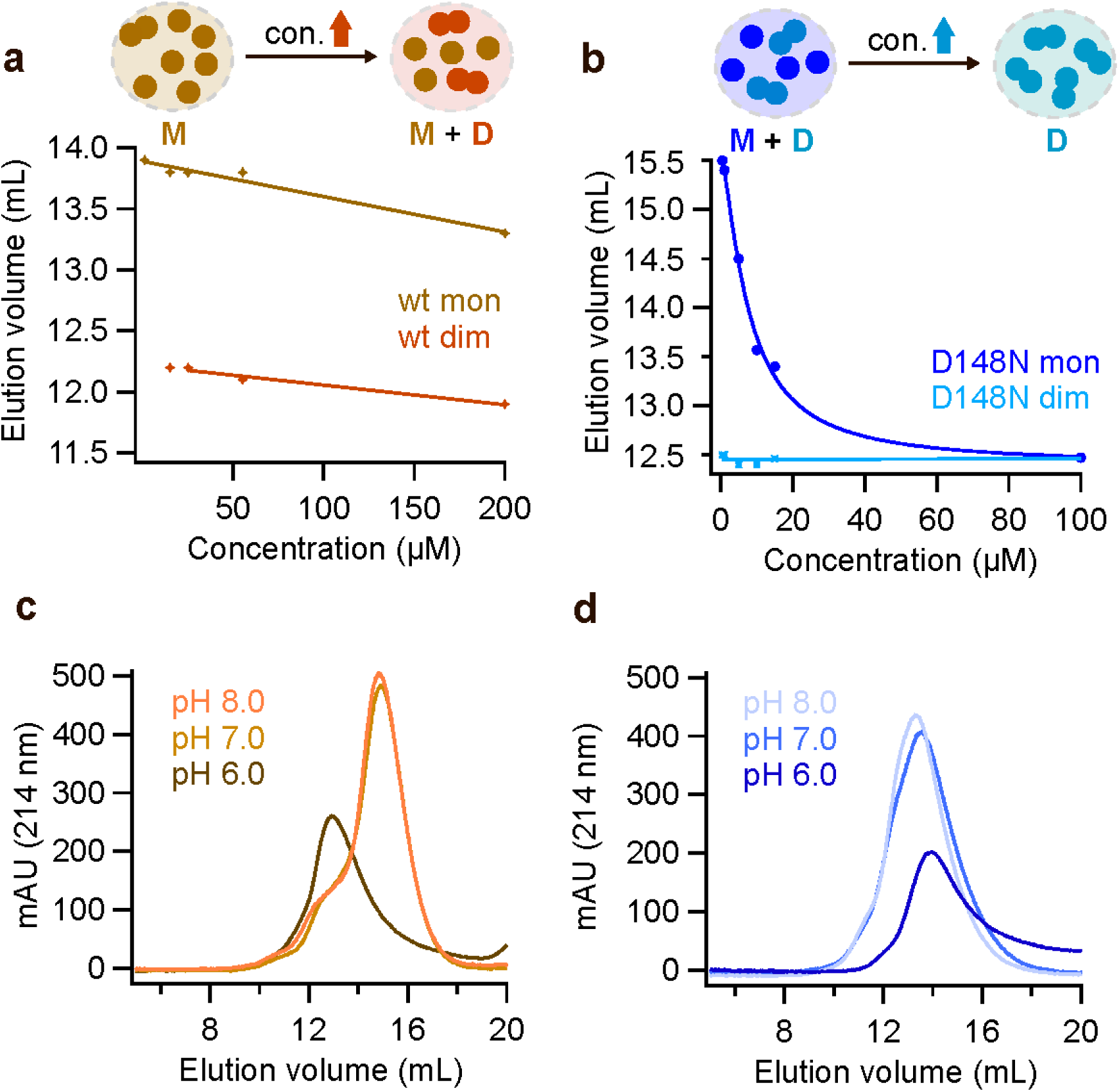
Concentration and pH dependent properties of rh BRICHOS from Bri2. (**a**) Concentration-dependent dimerization of rh wildtype Bri2 BRICHOS monomers, the details are shown in Supplementary Figure 7. The schematic inset shows that at low concentration the rh wildtype Bri2 BRICHOS monomer fraction is purely monomeric, while with increasing concentrations dimers form. M, monomer; D, dimer. (**b**) Concentration-dependent dimerization of rh Bri2 BRICHOS D148N monomers, the details are shown in Supplementary Figure 8. The schematic inset shows the presence of rh Bri2 BRICHOS D148N monomers and dimers already at low concentration, while at increased concentrations all monomers form dimers. M, monomer; D, dimer. pH dependent dimerization of (**c**) 10 µmol L^-1^ rh wildtype Bri2 BRICHOS monomers and (**d**) rh Bri2 BRICHOS D148N monomers analysed by SEC.

**Fig. S7.**
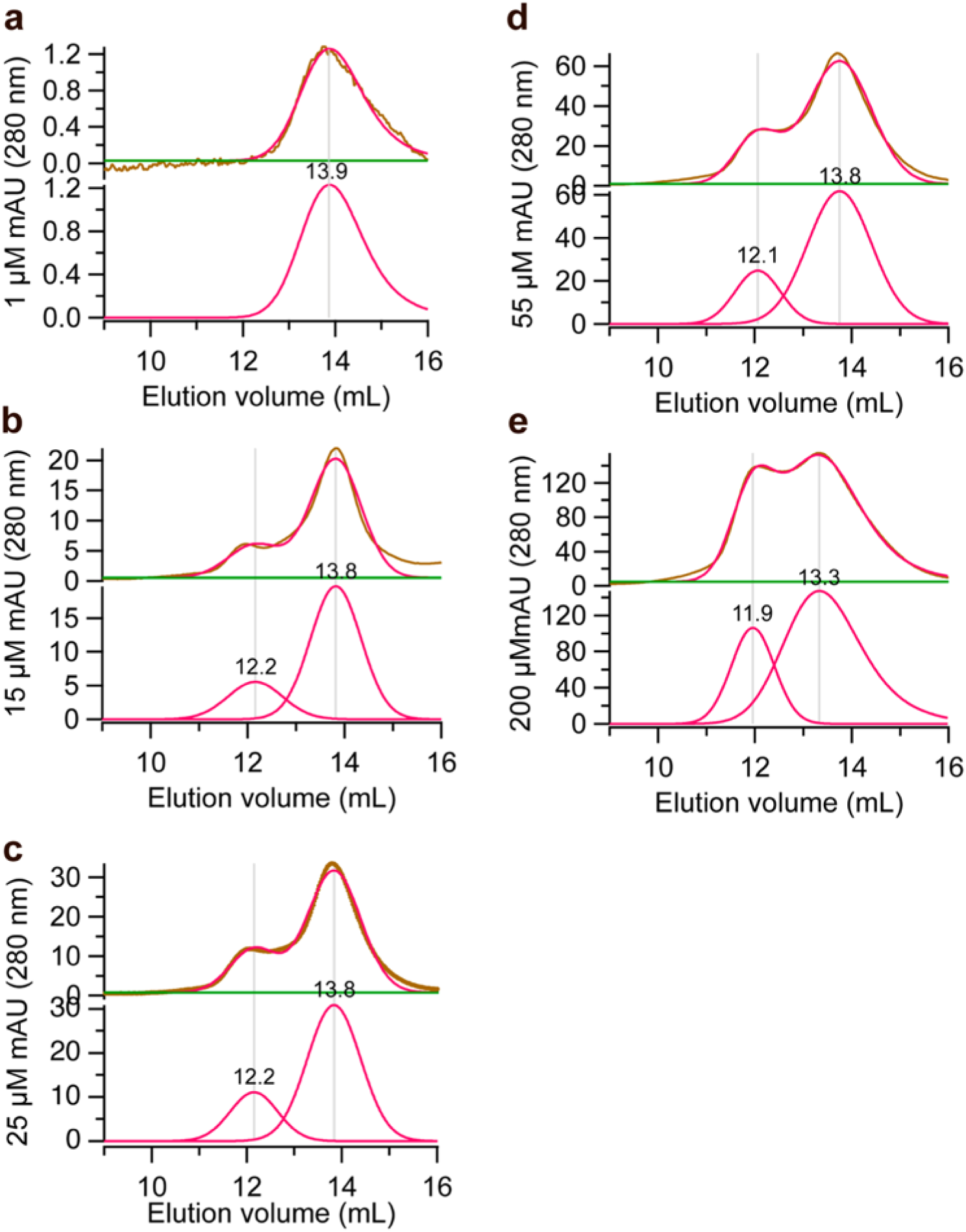
Rh wildtype Bri2 BRICHOS monomer concentration-dependent dimerization. SEC analysis of rh wildtype Bri2 BRICHOS monomer at concentrations of 1 (**a**), 15 (**b**), 25 (**c**), 55 (**d**) and 200 µmol L^-1^ (**e**). The top panel of each sub-figure is the raw SEC data (yellow) with gaussian fitted (red). The lower panel of each sub-figure is the multiple gaussian peak fitting (red) and the number on the top of each peak represents the elution volume.

**Fig. S8.**
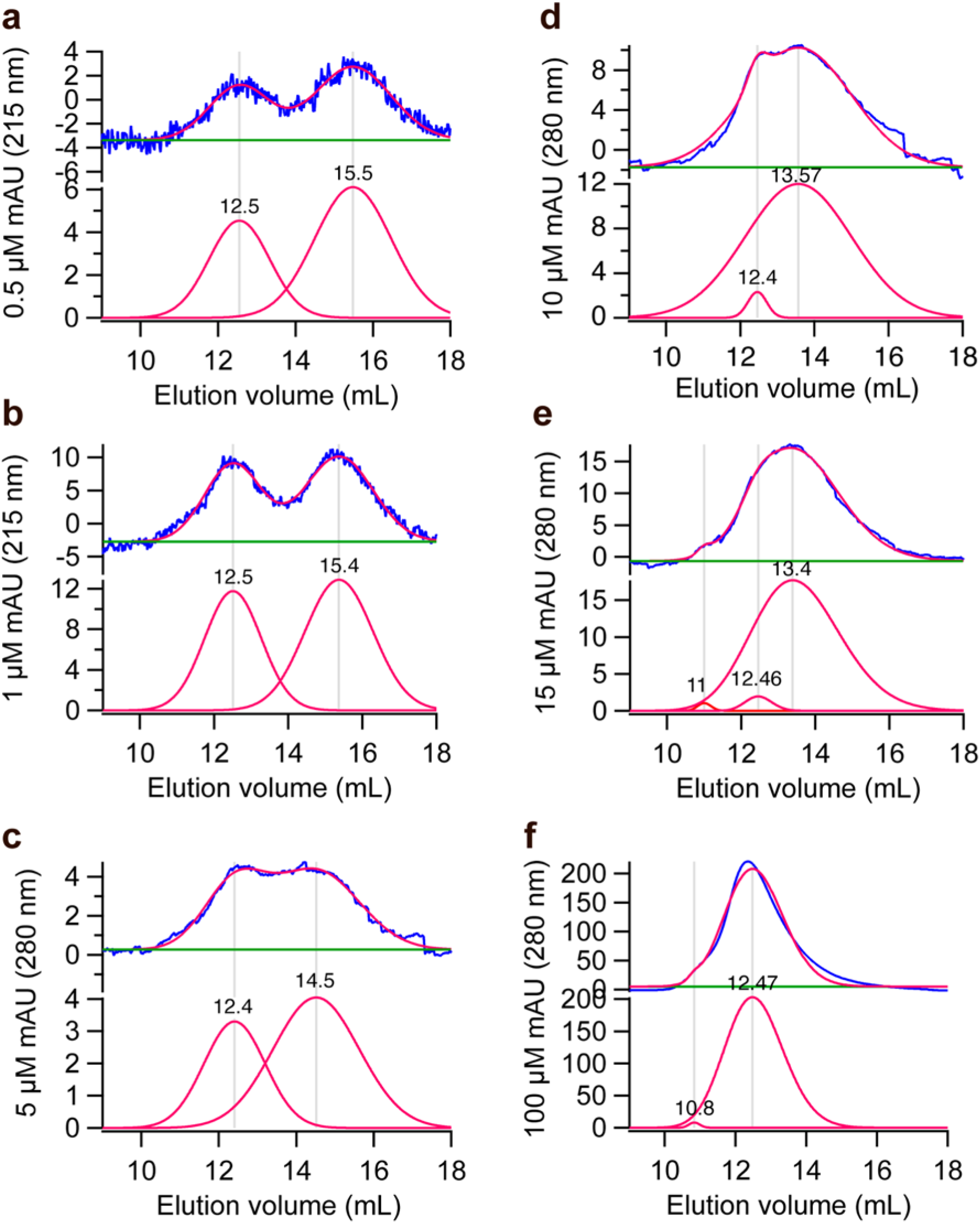
Rh Bri2 BRICHOS D148N monomer concentration-dependent dimerization. SEC analysis of rh Bri2 BRICHOS D148N monomer at concentrations of 0.5 (**a**), 1 (**b**), 5 (**c**), 10 (**d**), 15(**e**) and 100 µmol L^-1^ (**f**). The top panel of each sub-figure is the unprocessed data (yellow) with gaussian fitted (red). The lower panel of each sub-figure is the gaussian peak fitting (red) and the number on the top of each is the relevant elution volume.

**Fig. S9.**
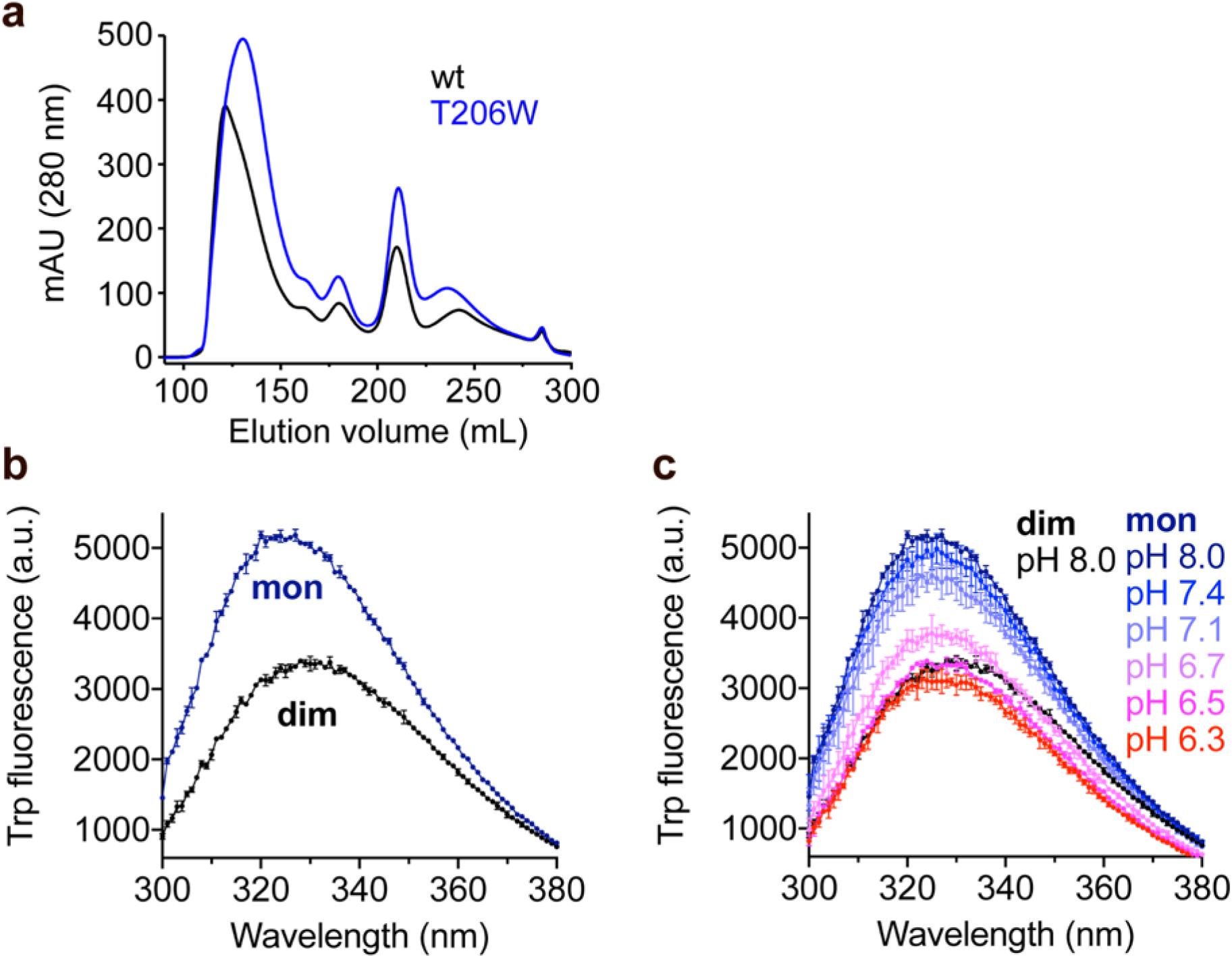
Trp fluorescence of rh Bri2 BRICHOS T206W mutant. (**a**) SEC of rh wildtype NT*-Bri2 BRICHOS and the rh NT*-Bri2 BRICHOS T206W mutant. (**b**) Rh Bri2 BRICHOS T206W monomers and dimers were diluted to 2 μmol L^-1^ in 20 mmol L^-1^ NaPi, 0.2 mmol L^-1^ EDTA at pH 8.0. (**b**) Rh Bri2 BRICHOS T206W monomers at different pHs from pH 6.3 to pH 8.0 were subjected to Trp fluorescence measurements, while the dimers were measured at pH 8.0. All the samples were prepared in duplicates, and excited at 280 nm (5 μm bandwidth) and fluorescence emission from 300–400 nm (10 μm bandwidth,1 nm step interval) was recorded. For the final fluorescence intensities, the results were corrected by subtracting the background fluorescence. The data are presented as means ± standard deviations.

**Fig. S10.**
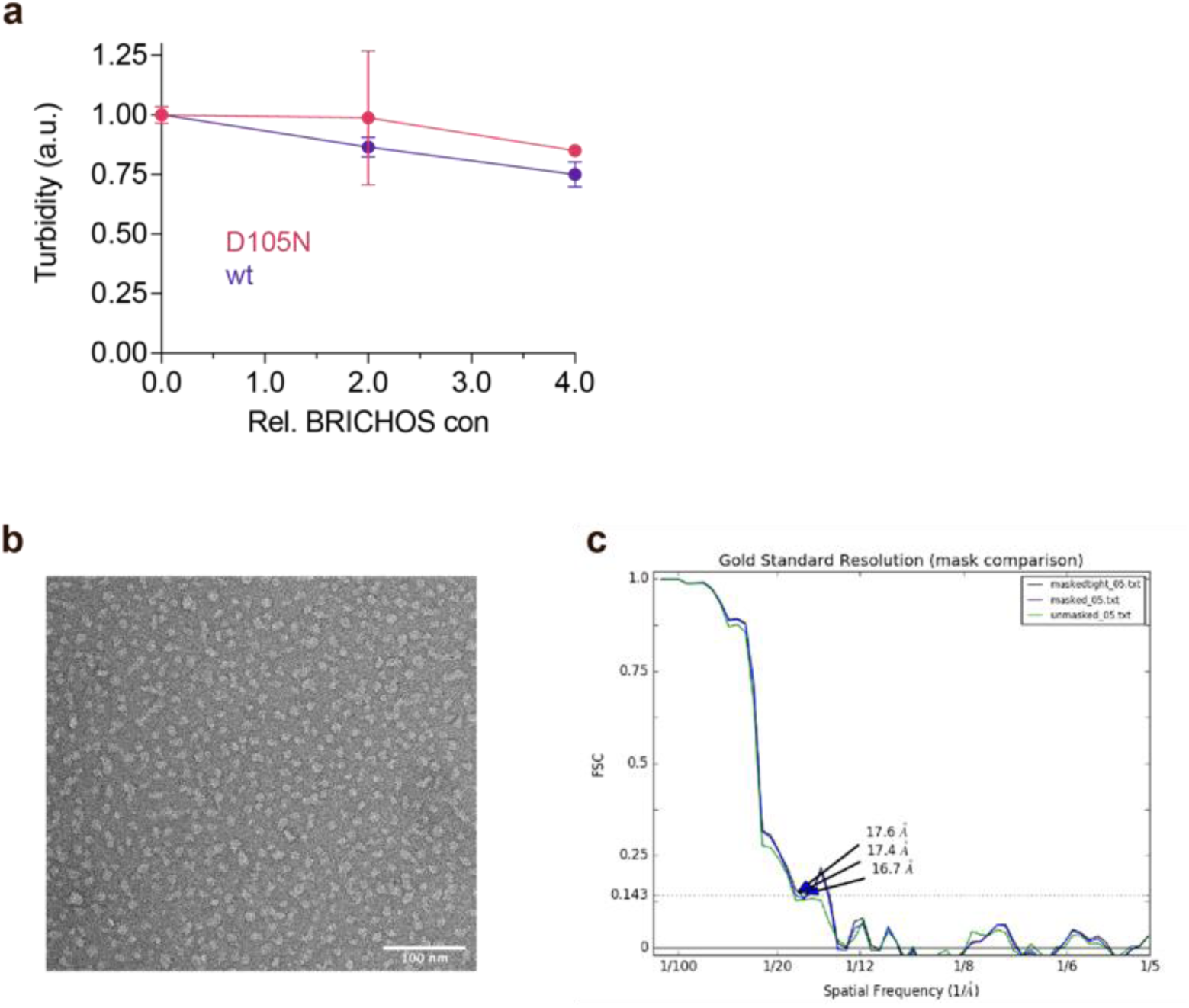
Electron microscopy analyses of rh Bri2 BRICHOS D148N oligomers. (**a**) Effects of rh proSP-C BRICHOS variants on CS aggregation at different molar ratios of BRICHOS:CS. The data are presented as means ± standard deviations. **(b**) Raw TEM micrograph of negatively stained rh Bri2 BRICHOS D148N oligomers. (**c**) The Fourier shell correlation (FSC) indicates a resolution of ∼17 Å for the reconstructed 3D density map determined at an FSC-value of 0.143 (dotted lines).

**Fig. S11.**
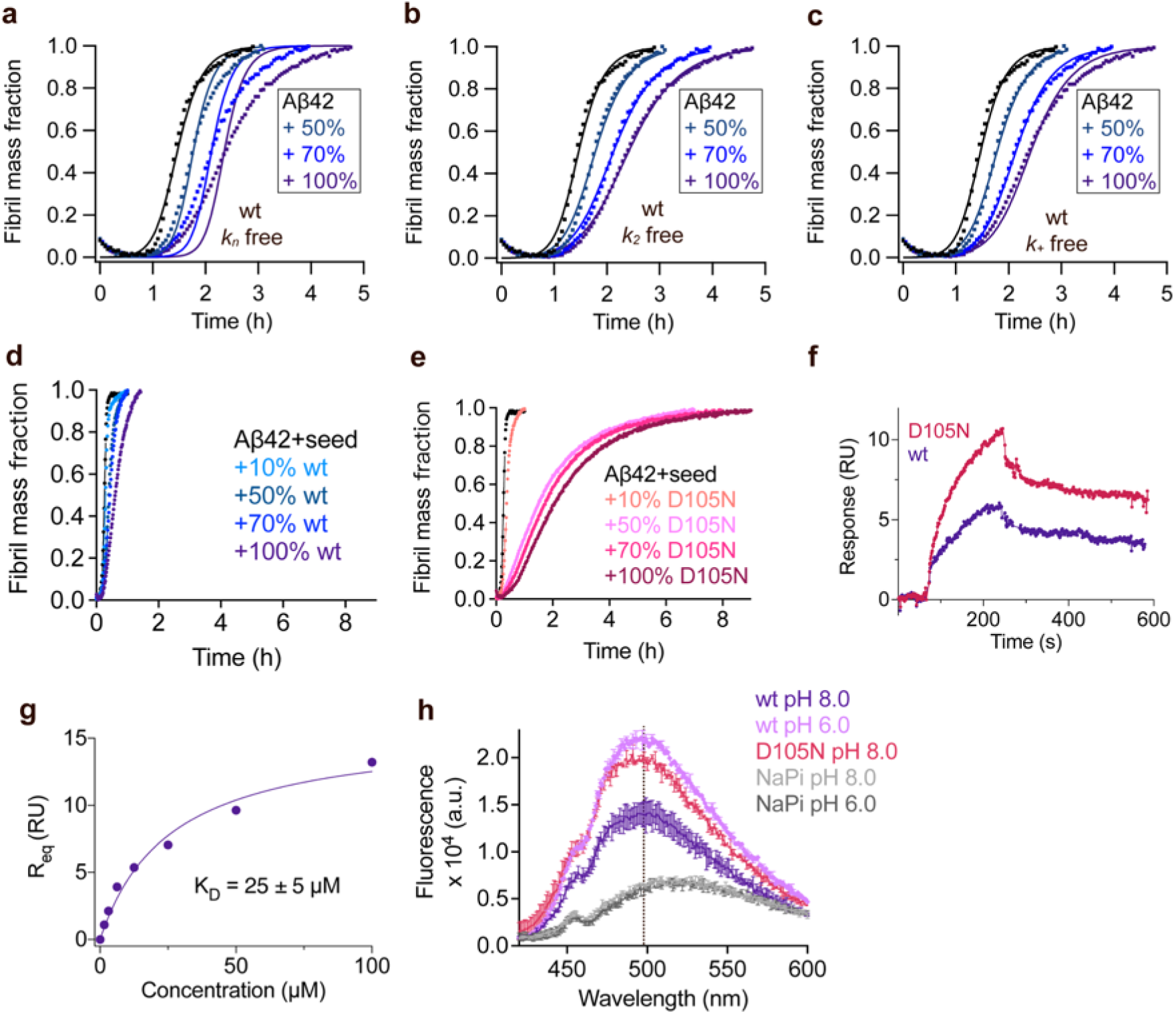
Rh wildtype proSP-C BRICHOS interferes with Aβ42 fibril formation. (**a–c**) Aggregation kinetics of 3 µmol L^-1^ Aβ42 in the presence of rh wildtype (wt) proSP-C BRICHOS at concentrations: 0 (black), 50 (cyan), 70 (blue), or 100% (purple) molar percentage referred to monomeric subunits relative to Aβ42 monomer concentration. The global fits (solid lines) of the aggregation traces (cross) were constrained such that only one single rate constant is the free fitting parameter. (**d** and **e**) Seeded aggregation traces of 3 µmol L^-1^ Aβ42 with 0.6 µmol L^-1^ preformed Aβ42 fibril seeds in the presence of different concentration of rh wildtype proSP-C BRICHOS (**d**) and the D105N mutant (**e**). (**f**) Analysis of rh wildtype (wt) proSP-C BRICHOS (purple) and the D105N mutant (red) binding to Aβ42 monomers detected by SPR. 25 µmol L^-1^ rh wildtype proSP-C BRICHOS and the D105N mutant were injected in over the chip surfaces, respectively. The response from the blank surface was subtracted from the immobilized surface response. (**g**) Steady state affinity for rh proSP-C BRICHOS D105N to immobilised Aβ42 monomers. Different concentrations of rh proSP-C BRICHOS D105N mutant, *i.e.* 0, 1.56, 3.13, 6.25, 12.5, 25, 50 and 100 µmol L^-1^, were individually injected over the chip surfaces. Steady state affinity was estimated by plotting the maximum binding responses versus BRICHOS concentrations. (**h**) Fluorescence emission of 2 µmol L^-1^ bis-ASN in sodium phosphate buffer at pH 8.0 (gray) or 6.0 (black) or in the presence of 1 µmol L^-1^ rh wildtype proSP-C BRICHOS at pH 8.0 or 6.0 and the D105N mutant.

**Fig. S12.**
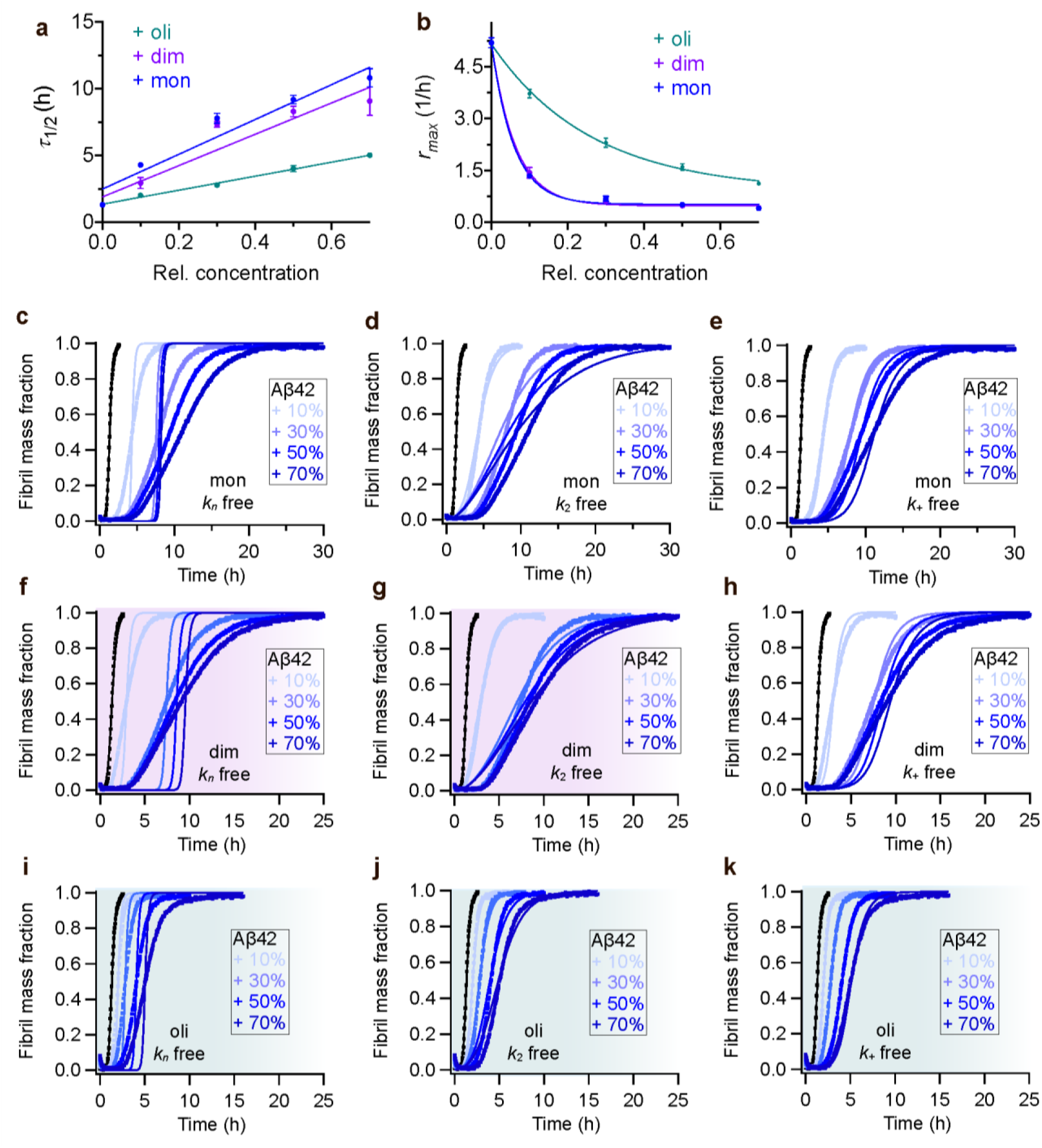
Rh Bri2 BRICHOS D148N inhibition of Aβ42 fibril formation. Values for *τ_1/2_* (**a**) and *r_max_* (**b**) extracted from the sigmoidal fitting of Aβ42 aggregation traces in the presence of different concentrations of rh Bri2 BRICHOS D148N species as shown in (**c–k**). (**c–e**) Aggregation kinetics of 3 µmol L^-1^ Aβ42 in the presence of rh Bri2 BRICHOS D148N monomers at different concentrations. The global fits (solid lines) of the aggregation traces (cross) were constrained such that only one single rate constant is the free fitting parameter, indicated in each panel. χ^2^ values describing the quality of the fits: 62 for *k_n_* free, 11.2 for *k*_2_ free and 3.7 for *k*_+_ free. (**f–h**) Aggregation kinetics of 3 µmol L^-1^ Aβ42 in the presence of rh Bri2 BRICHOS D148N dimers at different concentrations. The global fits (solid lines) of the aggregation traces (cross) were constrained such that only one single rate constant is the free fitting parameter, indicated in each panel. χ^2^ values describing the quality of the fits: 43 for *k_n_* free, 3.9 for *k*_2_ free and 7.8 for *k*_+_ free. (**i–k**) Aggregation kinetics of 3 µmol L^-1^ Aβ42 in the presence of rh Bri2 BRICHOS D148N oligomers at different concentrations. The global fits (solid lines) of the aggregation traces (cross) were constrained such that only one single rate constant is the free fitting parameter. χ^2^ values describing the quality of the fits: 8.6 for *k_n_* free, 1.4 for *k*_2_ free and 0.4 for *k*_+_ free. For the different D148N species, both *k*_2_ and *k*_+_ as sole free fitting rate, the fibrillization traces were described with similar χ^2^ values, suggesting both *k*_2_ and *k*_+_ might be affected, like the wildtype species.

## Supplementary Table

**Table S1. Species distribution and pairwise alignment of the multiple BRICHOS domains.** See separate file.

**Table S2. Summary of statistics performed in Fig. 3b and d**.

## Source data

**Figure 3-source data.**

**Figure 4-source data.**

**Figure 5-figure supplement 5-source data**

## Notes

### Competing Interest Statement

The authors have declared no competing interest.

